# Galectin-9 binding to HLA-DR in dendritic cells controls immune synapse formation and T cell proliferation

**DOI:** 10.1101/2024.11.29.625991

**Authors:** Andrea Rodgers Furones, Thijs Brands, Kristina Fedorova, Zacharias Wijfjes, René Classens, Lona Kroese, Martijn Verdoes, Guido van Mierlo, Annemiek B van Spriel, Laia Querol Cano

## Abstract

To initiate T cell-mediated immunity, dendritic cells (DCs) present antigens to specific T cells through the establishment of an immune synapse (IS). While the molecular mechanisms behind the formation of the IS on the T cell side are well understood, how IS components are organized at the DC membrane remain ill-defined. Galectins, a family of β-galactoside binding proteins, modulate immune cell function via the establishment of specific glycan-dependent or independent interactions. Nonetheless, the molecular mechanisms that underlie galectin function are poorly described and very little is known regarding their contribution to DC-mediated T cell activation. Here, we demonstrate that intracellular galectin-9 (gal9) in DCs is required for T cell activation. Murine and human DCs lacking gal9 showed impaired induction of CD4^+^, but not CD8^+^, T cell proliferation, suggesting a conserved function for gal9 in modulating DC–T cell interactions. Live-cell imaging revealed that galectin-9-depleted DCs fail to establish stable ISs with T cells, resulting in reduced T cell activation and proliferation. Unbiased co-immunoprecipitation and mass spectrometry identified HLA-II as a gal9 binding partner in DCs, and we observed a marked reduction of HLA-II recruitment to the immune synapse in DCs lacking gal9. Conditional gal9 knockout in DCs led to enhanced tumor growth *in vivo*, compared to their wild-type (WT) counterparts, underscoring a role for gal9 in T cell-dependent anti-tumor immunity. Collectively, this study provides the first detailed account of gal9-mediated HLA-II organization at the synaptic site of DCs, revealing a novel mechanism by which galectins orchestrate immune receptor positioning from within the cytoplasm to enhance CD4^+^ T cell activation.

## Introduction

Dendritic cells (DCs) are paramount to initiate cellular adaptive immunity due to their ability to recognise, digest and present antigens or tumor cells as peptides to T cells *via* the Major Histocompatibility complex (MHC) (Banchereau and Steinman, 1998; Steinman et al., 1997).

Antigen-specific interactions between DCs and T cells lead to the formation of the immunological synapse (IS), a highly ordered structure indispensable for T cell activation and function (Mempel et al., 2004). Assembly of an IS requires the recruitment of membrane receptors and signalling components to spatiotemporally organized domains at the interface between both cells (Friedl et al., 2005). Initial IS formation is dependent on the clustering of MHC and of the T cell receptor (TCR) (Dustin, 2014). This enables the recruitment of co-stimulatory molecules, integrins and other surface receptors that facilitate downstream signalling and dictate the strength and type of T cell immune response (de la Fuente et al., 2005; Friedl et al., 2005). The canonical organisation of the IS consists of a TCR-MHC rich central supramolecular activation cluster (c-SMAC), surrounded by a peripheral SMAC (p-SMAC) containing adhesion molecules (LFA-1/ICAM-1) that provide a mechanical scaffold for the IS and a distal SMAC (d-SMAC) that contains immunoinhibitory receptors such as the tyrosine phosphatase CD45 (Verboogen et al., 2016). Although the c-SMAC was initially thought to be the location of T cell signalling, it is now accepted that the c-SMAC poses a site of signalling termination and receptor recycling (Varma et al., 2006).

In contrast to the unifocal T cell-B cell synapse, the structure of the DC-T cell synapse appears to be multifocal, containing numerous segregated microdomains in which IS components are distributed at different points of the membrane, enabling multiple cell-cell interaction sites (Brossard et al., 2005; Dustin, 2014). The first phase of DC-T cell interactions consists of transient intermittent contacts that last approximately 10-12 min followed by a second phase, marked by the formation of stable DC clusters interacting with multiple T cells simultaneously, which can extend for about 2-3 h(Mempel et al., 2004; Shakhar et al., 2005). Upon DC-T cell disengagement, a third phase follows where T cells regain their motility and proliferate (Mempel et al., 2004). Importantly, DC-T cell interactions rely on the reorientation of the microtubule organizing centre (MTOC) and on cytoskeletal remodelling in DCs to drive the recruitment of IS components to specific areas within the contact zone and to promote DC polarization towards the interacting T cells (Boes et al., 2002; Compeer et al., 2014; Malinova et al., 2016). Nonetheless, while the molecular mechanisms governing IS membrane organization on the T cell side are well understood, the organization of the IS on the DC side remains ill-defined.

Galectins are a family of soluble β-galactoside binding proteins that share at least one conserved carbohydrate recognition domain (CRD). Through specific carbohydrate-dependent interactions on the cell surface galectins modulate receptor mobility, clustering or stability (Querol Cano et al., 2024). In addition, galectins exert intracellular functions through both glycan-independent and glycan-dependent interactions (Johannes et al., 2018; Nabi et al., 2015; Querol Cano et al., 2024). Galectins modulate T cell activation either through their interactions with cell surface glycans or by influencing T cell signalling. As key organizers of glycans, galectins contribute to the formation of structured lattices on the cell surface, such as the N-glycosylated TCR lattice in T cells, which prevents unintended activation in resting cells (Demetriou et al., 2001). Extracellular galectin-3 inhibits T cell activation by restricting the lateral movement of the TCR within the IS and dissociating CD8 from the TCR (Demotte et al., 2008). Additionally, galectin-3 influences synapse dynamics by sequestering LFA-1 in membrane structures, thereby affecting LFA-1 recruitment and activation (Demotte et al., 2008; Petit et al., 2016). Illustrating the pleiotropic functions of galectins in T cells, intracellular galectin-9 (gal9) enhances T cell activation by supporting TCR-CD3 complex formation and signaling pathways (Chen et al., 2020; Lhuillier et al., 2015) whereas galectin-3 triggers TCR internalisation effectively inhibiting T cell activation (Chen et al., 2009). Despite these well-characterized functions on T cells, the role of galectins at the DC side of the IS remains unexplored.

Here, we demonstrate that DCs require gal9 to interact with CD4^+^ T cells. We identified an interaction between gal9 and HLA-DR crucial for its recruitment to the contact zone between DC and T cells and subsequent T cell activation and proliferation. Our work reveals a novel function for intracellular gal9 in regulating DC-IS formation, important for T cell activation *in vivo*.

### Materials and methods Generation of monocyte-derived dendritic cells

Dendritic cells were derived from peripheral blood monocytes isolated from a buffy coat (Sanquin, Nijmegen, The Netherlands) as previously described(Querol Cano et al., 2019). Monocytes isolated from healthy blood donors (informed consent obtained) were cultured for up to five days in RPMI 1640 medium (Life Technologies, Bleiswijk, Netherlands) containing 10 % foetal bovine serum (FBS, Greiner Bio-one, Alphen aan den Rijn, Netherlands), 1 mM ultra-glutamine (BioWhittaker), antibiotics (100 U/ml penicillin, 100 µg/ml streptomycin and 0.25 µg/ml amphotericin B, Life Technologies), IL-4 (500 U/ml, Miltenyi Biotec) and GM-CSF (800 U/ml, #130-093-868, MiltenyiBiotec) in a humidified, 5 % CO_2_. On day 3, medium was refreshed with new IL-4 (500 U/ml, Miltenyi Biotec) and GM-CSF (800 U/ml, Miltenyi Biotec). On day 6, moDCs were supplemented with a maturation cocktail: IL-6 (15 ng/ml, #130-093-933, Miltenyi Biotec), TNF-α (10 ng/mg, #130-094-014 Miltenyi Biotec), IL-1β (5 ng/ml, #130-093-898 Miltenyi Biotec) and PGE2 (10 ug/ml, Pfizer). When necessary, moDCs were treated with recombinant galectin-9 protein (AF2045, R&D systems) at a final concentration of 1 µg/ml or with an anti-galectin-9 blocking antibody (# MABT834, Sigma-Aldrich) at a final concentration of 10 µg/ml.

### Mice

The conditional Lgals9 mouse strain (MGI:6466637) was generated on the C57Bl/6JRj background using pronuclear microinjection in mouse zygotes. The injection mixture consisted of water with 200ng/µl Cas9 protein (IDT), two sgRNA (25ng/µl each) targeting the intronic sequence flanking exon 3 of the *Lgals9* gene (5’-CGGGACTAGAGCGTGTCTTA GGG-3’ and 5’-TAAAACCCAGCGGGCGAATG GGG-3’) and 15ng/µl of a long single stranded DNA oligo containing exon 3 of Lgals9 flanked by two loxP recombination sites and homology arms. The hom. floxed mice were sequence verified.

Sex- and age-matched C57Bl/6J WT (*lgals9^+/+^*), and *lgals9^-/-^* littermate mice were bred at the Central Animal Laboratory in Nijmegen, the Netherlands. Both male and female age-matched mice were used across replicate experiments. OT-I and OT-II mice were sourced from Charles River. All mice were housed in top-filter cages, provided a standard diet, and had unrestricted access to water and food. Mice were used at ages ranging from 6 to 18 weeks. All murine studies complied with European legislation (directive 2010/63/EU of the European Commission) and were approved by local authorities (CCD, The Hague, the Netherlands) for the care and use of animals with related codes of practice.

### Isolation and culture of primary cells

Human pan T cells and Pan naïve T cells were isolated from peripheral blood mononuclear cells derived from healthy individuals (Sanquin, Nijmegen, the Netherlands) using the Pan T cell isolation kit (130-096-535, Miltenyi Biotec) and the Pan Naïve T cell kit (130-097-095, Miltenyi Biotec) according to manufacturer’s instructions. After isolation, fresh T cells were cultured in X-VIVO-15 (Lonza) supplemented with 2 % human serum (HS, Sigma-Aldrich).

For murine DCs, lymph nodes and spleens from wild-type C57BL/6J, *lgals9^-/-^*, *cd11c^cre^lgals^fl/fl^* mice were isolated and meshed to obtain single cell suspension. Cells were meshed through a 100 μm cell strainer by using a syringe plunger. Cell suspension was spun at 400xg for 5 min and resuspended in 2 ml of 1x ammonium chloride potassium (ACK) solution for the lysis of erythrocytes. After 5 min of incubation at room temperature (RT) cells were washed with 20 ml of PBS 2 times. DCs were isolated using the CD11c MicroBeads UltraPure isolation kit (#130-125-835, Miltenyi Biotec) and immediately used for functional studies.

### Small interfering RNA knockdown

On day 3 of DC differentiation, cells were harvested and subjected to electroporation. Three custom stealth small interfering RNA (siRNA) were used to silence galectin-9 (LGALS9HSS142807, LGALS9HSS142808 and LGALS9HSS142809) (Invitrogen). Equal amounts of the siRNA ON-TARGETplus non-targeting (NT, here referred as wildtype, WT) siRNA#1 (Thermo Scientific) were used as control. Cells were washed twice in PBS and once in OptiMEM without phenol red (Invitrogen). A total of 15 μg siRNA (5 μg from each siRNA) was transferred to a 4-mm cuvette (Bio-Rad) and 5-10x10^6^ DCs were added in 200 μl OptiMEM and incubated for 3 min before being pulsed with an exponential decay pulse at 300 V, 150 mF, in a Genepulser Xcell (Bio-Rad, Veenendaal, Netherlands), as previously described (Querol Cano et al., 2019; Santalla Mendez et al., 2023). Immediately after electroporation, cells were transferred to preheated (37 °C) phenol red–free RPMI 1640 culture medium supplemented with 1 % ultraglutamine, 10 % (v/v) FCS, IL-4 (300 U/ml), and GM-CSF (450 U/ml) and seeded at a final density of 5x10^5^ cells/ml.

### Flow cytometry

To determine depletion of galectin-9 following siRNA transfection, human single cell suspensions were stained with a goat anti-galectin-9 antibody (AF2045, R&D systems) at 8 μg/ml for 30 min at 4 °C. Before staining, moDCs were incubated with 2 % human serum for 10 min on ice to block non-specific interaction of the antibodies with Fc receptors. A donkey-anti goat secondary antibody conjugated to Alexa Fluor 488 was used (Invitrogen; 1:400 (v/v)).

moDCs were incubated for 30 min on ice with the antibodies (see Table 1). All antibodies were used within a range of a final 1:25 to 1:50 (v/v) dilution in cold PBS containing 0.1 % BSA, 0.01 % NaH_3_ (PBA) supplemented with 2 % HS.

**Table 1.**

| Application | Antibody Target | Company | Catalog Number | Clone | Color | Dilution |
| --- | --- | --- | --- | --- | --- | --- |
| Human DC maturation | HLA-DR | BD BioSciences | # 555811 | G46-6 | FITC | 1:50 |
|  | HLA-DR | BioLegend | # 307646 | L243 | BV-510 | 1:50 |
|  | CD40 | BD BioSciences | # 561215 | 5C3 | PE-Cy7 | 1:50 |
|  | CD80 | BD BioSciences | # 557227 | L307.4 | PE | 1:50 |
|  | CD80 | BD BioSciences | # 561135 | L307.4 | PE-Cy7 | 1:50 |
|  | CD86 | BD BioSciences | # 555660 | 2331 | APC | 1:50 |
|  | CD11b | BioLegend | # 301342 | ICRF44 | APC-Cy7 | 1:50 |
|  | HLA-ABC | BD BioSciences | # 555553 | G46-2.6 | PE | 1:50 |
|  | HLA-ABC | Miltenyi | # 130-101-466 | REA230 | APC | 1:50 |
|  | PDL-1 | BD BioSciences | # 563738 | MIH1 | BV-421 | 1:50 |
|  | CD70 | BD BioSciences | # 555835 | Ki-24 | PE | 1:50 |
|  | CD14 | BD BioSciences | # 560180 | MoP9 | APC-H7 | 1:50 |
| Human T cell proliferation | CD25 | BD BioSciences | # 555434 | M-A251 | APC | 1:25 |
|  | CD69 | BD BioSciences | # 555530 | FN50 | FITC | 1:25 |
|  | CD8 | BioLegend | # 344708 | SK1 | PerCP | 1:25 |
|  | CD3 | eBioscience | # 25-0038-42 | UCHT1 | PE-Cy7 | 1:25 |
|  | CD4 | BD BioSciences | # 560158 | RPA-T4 | APC-H7 | 1:25 |
| Human T cell exhaustion | CD4 | eBioscience | # 25-0047-42 | SK3 | PE-Cy7 | 1:50 |
|  | CD8 | BD Biosciences | # 555366 | RPA-T8 | FITC | 1:100 |
|  | TIGIT | BD Biosciences | # 747846 | 741182 | BB700 | 1:20 |
|  | Tim-3 | BioLegend | # 345012 | F38-2E2 | APC | 1:100 |
|  | PD1 | BD Biosciences | # 563076 | EH12.1 | BV510 | 1:15 |
| Human CD4+- cytokine production | IL-10 | BioLegend | # 501420 | JES3-9D7 | PE-Cy7 | 1:20 |
|  | IL-4 | Miltenyi Biotec | # 130-123-698 | 7A3-3 | PE | 1:25 |
|  | IFN-γ | Miltenyi Biotec | # 130-113-497 | REA600 | FITC | 1:50 |
|  | IL-17A | BioLegend | # 512310 | BL168 | af647 | 1:25 |
|  | CD4 | BioLegend | # 300528 | RPA-T4 | PerCP | 1:25 |
| Human CD8+- granular production | Granzyme-B | BioLegend | # 372208 | QA16A02 | PE | 1:30 |
|  | Perforin | BioLegend | # 308110 | dG9 | af647 | 1:30 |
|  | CD8 | BD Biosciences | # 555366 | RPA-T8 | FITC | 1:100 |
| Human T regulatory phenotype | CD4 | eBioscience | # 25-0047-42 | SK3 | PE-Cy7 | 1:50 |
|  | CD25 | BD Biosciences | # 555434 | M-A251 | APC | 1:100 |
|  | Foxp3 | eBioscience | # 53-4776-42 | PCH101 | af488 | 1:50 |
|  | CD127 | eBioscience | # 12-1278-42 | eBioRDR5 | PE | 1:100 |
| Human CD4+<br>T helper<br>subsets | CD4 | eBioscience | # 12-1278-42 | eBioRDR5 | PE | 1:50 |
|  | GATA3 | BD Biosciences | # 560078 | L50-823 (RUO) | af647 | 1:100 |
|  | T-bet | Invitrogen | # 45-5825-82 | 4B10 | PerCP-Cy5.5 | 1:100 |
|  | RORγt | Invitrogen | # 12-6988-82 | AFKJS-9 | PE | 1:200 |
| Mouse<br>dendritic cell<br>panel | XCR1 | Biolegend | # 148216 | ZET | BV421 | 1:50 |
|  | CD11c | Biolegend | # 117353 | N418 | BV510 | 1:50 |
|  | CD172a | Invitrogen | # 17-1721-82 | P84 | BV605 | 1:50 |
|  | CD3 | Biolegend | # 100204 | 17A2 | FITC | 1:50 |
|  | CD4 | Biolegend | # 100510 | RM4-5 | FITC | 1:50 |
|  | Ly6G | Biolegend | # 127606 | 1A8 | FITC | 1:50 |
|  | Gal9 | Biolegend | # 136101 | RG9-35 | PE | 1:25 |
|  | MHCII | Biolegend | # 107624 | M5/114.15.2 | PerCP | 1:50 |
|  | CD103 | Biolegend | # 121426 | 2E7 | PeCy7 | 1:50 |
|  | SigleCH | eBioscience | # 51-0333-82 | eBio440c | af647 | 1:50 |
| Mouse<br>myeloid panel | F4/80 | Biolegend | # 123131 | BM8 | BV421 | 1:50 |
|  | CD11c | Biolegend | # 117353 | N418 | BV510 | 1:50 |
|  | CD11b | BD | # 553309 | M1/70 | BV605 | 1:50 |
|  | Ly-6C | BD | # 553104 | AL-21 | FITC | 1:50 |
|  | Gal9 | Biolegend | # 136101 | RG9-35 | PE | 1:25 |
|  | MHCII | Biolegend | # 107624 | M5/114.15.2 | PerCP | 1:50 |
|  | Ly-6G | Biolegend | # 127618 | 1A8 | PeCy7 | 1:50 |
|  | CD3 | Biolegend | # 100312 | 145-2C11 | APC | 1:50 |
|  | CD4 | Biolegend | # 100516 | RM4-5 | APC | 1:50 |

To determine T cell activation or proliferation, harvested cells at different timepoints were blocked in PBA buffer supplemented with 2 % HS for 10 min. Afterwards, T cells were stained in 2 % HS PBA buffer with the antibodies (see Table 1) at 1:25 (v/v) dilution for 20 min.

To phenotype expanded T cells, harvested cells were divided into different flowcytometry panels. Exhausted T cells were blocked as aforementioned and stained with the antibodies (see Table 1). CD4+ and CD8+ intracellular cytokines were evaluated by first stimulating with PMA (25 ng/ml, Calbiochem), Ionomycin (0.5 μg/ml, Sigma-Aldrich; # I0634) and Brefeldin A (10 ng/ml, Cayman chemicals). After 4 h of treatment at 37°C, cells were washed and stained with the eFluor780 viability dye (1:2000) as aforementioned. Next, cells were fixed and permeabilised using the BD Cytofix/Cytoperm™ Fixation/Permeabilization Kit (BD Biosciences; # 554714) following the manufacturer’s instructions. After blocking, cells were stained with the antibodies (see Table 1). Other panels phenotyping fixed, permeabilised using the FOXP3 transcription factor staining kit (00-5523-00, ThermoFisher) according to the manufacturer’s instructions and blocked as mentioned above. To detect T regulatory T cell phenotypes, expanded T cells were stained with the antibodies (see Table 1). T cell-helper subsets were identified by staining with the antibodies (see Table 1).

Splenic and lymph node-cell suspensions from C57Bl/6J WT (*lgals9^+/+^*), and *lgals9^-/-^* and *cd11c^cre^lgals^fl/fl^*animals were blocked with 5 % human serum for 10 min on ice. Cells were equally splitted into two different stainings (dendritic cell subset and myeloid panels) before incubating with the antibodies (see Table 1, 30 min at 4 °C). Cells were then washed two times and incubated with Brilliant Violet 605™ Streptavidin (Biolegend, #405229, at 1:100 (v/v)) for 15 min at 4 °C in the dark.

Before measuring, live/dead staining was performed with eFluor780 or zombie violet viability dyes (1:2000, ThermoFisher) in PBS for 20 min at 4°C. If not previously, cells were fixed with a 4% PFA solution 10 min at RT. After 2-3 washes, stained cells were analyzed by flow cytometry using a FACSVerse or a FACSLyric instrument (BD, Frankllin lakes, NJ, USA) and later analyzed using the Cytobank platform (Beckman Coulter Life Sciences).

### Mixed lymphocyte reaction (MLR)

Allogeneic Pan T cells were stained with CFSE or CellTrace™ Violet Cell Proliferation Kit (C34554 or C34557 ThermoFisher), 5 μM; Life Technologies) for 30 min at 37°C; thereafter, reaction was stopped by blocking with 1:1 (v/v) FCS 5-10 min at RT. Cells were washed extensively and counted. WT or galectin-9 knockdown DCs were cultured with CTV/CFSE-labelled T cells in a ratio 1:10 for 6 days at 37°C, 5% CO_2_. After the 6 days, T cells were measured on the flow cytometer to evaluate the loss of cell trace dye expression. When needed, galectin-9 knocked down DCs were incubated with 1 µg/ml human recombinant galectin-9 protein (R&D systems, #2045-GA-050) for 30 min prior to being co-cultured with T cells. Alternatively, non-targeting cells were treated with 10 µg/ml galectin-9 blocking monoclonal antibody (Merck Millipore, # MATB834, clone 9S2).

The proliferation index (PI), was used to quantify the number of times an original T-cell divided ((PI)/2), where PI was the percentage of cells in each peak of division). Additionally, the fold change was calculated as [% proliferating T cells in stimulated condition]/[% proliferating cells in untreated conditions].

### Naïve T cell expansion and autologous T cell activation assays

WT or gal-9-depleted moDCs and allogeneic naïve T cells were co-cultured at a ratio of 1:5 (DC:T cell) in a 96 U bottom culture plate. After day 6 (and periodically every two days until day 13-14 of culture), 20 U/ml of recombinant IL-2 (130-097-744, Miltenyi Biotec) was added to the co-cultures. Resting T cells were counted and plated for phenotypical analysis.

To assess MHC-II-mediated antigen presentation, day 6 WT or galectin-9 depleted moDCs were cultured with 0.5 μg/ml tetanus toxoid (TT) (Sigma Aldrich; #582231) for 2 h. Loaded DCs were washed twice with PBS and co-cultured with autologous CTV-labelled CD4^+^ T cells in a 96-round well microplate at a ratio of 1:50 (DC:T cell) for 6 days prior to cells being harvested and T cell proliferation determined by flow cytometry.

To assess MHC-I-mediated antigen presentation on day 6, HLA-A*0201-type WT or galectin-9 knockdown moDCs were seeded at a density of 1x10^6 cells/ml and treated with 1 μg/mL LPS (Invivogen, # vac-3pelps) together with 1 μM of gp-100 peptide (custom synthesized by Genscript) or 1 μM (irrelevant) NY-ESO1 peptide (SLLMWITQC; custom synthesized by Genscript) for 2 h at 37 °C, 5% CO_2_. After this time, cells were washed and seeded at the aforementioned density in X-VIVO media supplemented with 2% human serum for 2 h at 37 °C, 5% CO_2_. Autologous CD8^+^ T cells were washed in PBS, resuspended in 250 µL red phenol-free X-VIVO medium and transferred into a 4-mm cuvette (Bio-Rad). Directly prior to electroporation, 20 μg/10x10^6 cells HLA-A2-gp100 TCR mRNA (provided by BioNtech) was added into the cell suspension. Cells were then pulsed with a square wave protocol at 500 V, 3 ms, 1 pulse, in a Genepulser Xcell (Bio-Rad, Veenendaal, The Netherlands) and transferred for 1 h (37°C, 5% CO2) into 1 mL red phenol-free X-VIVO medium (supplemented with 5% human serum without antibiotics). moDCs were co-cultured with autologous transfected T cells at a 1:5 ratio (DC:T cell) in 96 well round bottom plates for 3 days prior to cells being harvested and T cell proliferation determined by flow cytometry. The expression of the gp100-specific TCR was confirmed using an MHC-Dextramer (HLA-A*0201/YLEPGPVTA) (Immudex) (Fig. S3B).

Splenic OT-I or OT-II mouse T cells were isolated with Pan T Cell Isolation Kit II, mouse isolation kit (Miltenyi). Wild type or *galectin-9* ^-/-^ CD11c+ cells were treated for 2 h with 1 μg/mL LPS and primed with 25 µg/mL or 75 µg/mL of OVA EndoFit™ (InvivoGen, # vac-pova) or 1 µg/mL OVA peptides (OVA_257-264_ for OT-I cells or OVA_323-339_ for OT-II cells, from InvivoGen, #vac-sin and #vac-isq, respectively). Primed DCs and OT-I or OT-II cells were co-cultured at a ratio of 1:5 for 3 days after which T cell proliferation was determined as before. When necessary, *galectin-9* ^-/-^ DCs were incubated with 2 µg/ml mouse recombinant galectin-9 protein (R&D systems, #3535-GA-050) for 1 h prior to being co-cultured with T cells.

### ELISA

Enzyme-linked immunosorbent assay kits (Invitrogen, ThermoFisher Scientific) were used to measure IFN-γ (KIT REF: 88.7316.88) from supernatants obtained from DC-T cell co-cultures on day 6 and from 48, 72, 96 and 120 h. Supernatants were diluted 1:50 (for the 6-day co-cultures) or 1:2 (for the 48-120 h co-cultures). Protocols were performed following manufactures instructions. Standard curves were run at the same time and used to calculate the concentration of cytokines in the samples.

### Immunofluorescence and confocal microscopy

On day 6, moDCs were treated with 1 µg/mL Staphylococcal enterotoxin B or super antigen B (S4881, Sigma Aldrich) for 1 h at 37°C (5% CO_2_). After extensive washing, 60.000 mature wild type or galectin-9 depleted moDCs were incubated in 96-well low attachment plates with autologous T cells (1:2 ratio) for 2 h prior to being transferred to a 12 mm PLL-coated coverslip (P4707-50 ml, Sigma-Aldrich). Cells were left to adhere for 10 min prior to being fixed in freshly prepared 4% paraformaldehyde (PFA) for 10 min at RT and subsequently incubated for 20 min in 0.1 M quenching solution (NH_4_Cl in PBS). Coverslips were then washed twice with PBS and permeabilized in permeabilization buffer (2.5% donkey serum and 0.1% saponin) for 30 min prior to being incubated o/n and at 4 °C with specific antibodies against galectin-9 (R&D biosystems, #AF2425, 1:40 final dilution). The following day, coverslips were incubated for 1 h in permeabilisation buffer at RT, in the dark with the following secondary and conjugated antibodies: donkey-anti-goat alexa 568 (ThermoFisher Scientific, #A11055) at 1:400 (v/v) dilution, conjugated antibodies: anti-human TCR alpha (α) and beta (β) FITC (488) (BD Biosciences, 43720) at 1:50 (v/v) dilution, anti-human HLA-DR, DP, DQ Alexa flour 647 (BD Biosciences, 563591, clone Tü39) at 1:50 (v/v) dilution. After incubation, cells were washed thoroughly with PBS, incubated with 0.3 ug/ml 4′-6-diamidino-2-phenylindole (DAPI 1:3000 dilution) for nuclear staining, washed again with PBS and embedded in glass slides using 8 µl Mowiol (Calbiochem). Coverslips were allowed to dry overnight at RT in the dark and stored at 4 °C until imaging. Samples were imaged with both a Leica DMI6000 epi-fluorescence microscope fitted with a 63 × 1.4 NA oil immersion objective, a metal halide EL6000 lamp for excitation, a DFC365FX CCD camera and GFP and DsRed filter sets (all from Leica, Wetzlar, Germany) as well as a Zeiss LSM900 confocal laser scanning microscope equipped with the Airyscan module. For the images acquired with the Leica DMI6000 epi-fluorescence microscope we used a 63 × 1.4 NA oil immersion objective, a metal halide EL6000 lamp for excitation, a DFC365FX CCD camera and GFP and DsRed filter sets (all from Leica, Wetzlar, Germany). Focus was kept stable with the adaptive focus control from Leica. For the images acquired with eh Zeiss LSM900 microscope we used a 63 × EC Epiplan-NEOFLOUAR oil immersion objective and three laser module URGB (405, 488, 561, 640 nm). Images were analyzed with ImageJ software. Fluorescence overlap was quantified using a custom-made macro in Fiji ImageJ. The Mander’s and Pearson’s correlation coefficients were calculated using the JACoP plugin and the line scan graphs representing the fluorescence cross-sections were quantified using the Plots profile command in Fiji ImageJ and depicted using GraphPad Prism 8 or 10 software. The cross-sections were established by drawing a perpendicular line from the center of the immune synapse towards the plasma membrane at the rear of the cell.

### Live cell microscopy

Day 6 moDCs were treated with super antigen B as previously stated. Before co-culturing and to enable cell segmentation during data analysis, DCs were stained with FarRed Cell Trace^TM^ dye (C34564, Thermo Fisher) according to manufacturer’s instructions and autologous T cells were stained with CFSE Cell Trace^TM^ dye (C34554, Thermo Fisher). Both cell types were seeded on a black 96 well, f-bottom (chimney well) microplate (655076, Greiner) and immediately acquired.

Time-lapsed video microscopy was performed using the Celldiscoverer7 (Zeiss), using the 20x objective with 0.5x tube lens or the BD Pathway 855 spinning disk confocal microscope (BD Bioscience), the atto vision software (BD Bioscience) and the 10X objective (Olympus). Sequential images were acquired every 3 min for 2 h using the 555/30 excitation filter (Chroma), 631/33 excitation filter (Chroma), emission filter 84101 (Chroma) and dichroic filter 84000 (Chroma). Time-lapse sequences were analyzed with the Trak Mate plugin (Fiji) to measure cell velocity.

### RNAseq

Total RNA was extracted from mature moDC using zymo quick-RNA miniprep kit R1055 according to manufacturer instructions and used as input for library preparation using KAPA RNA HyperPrep Kit (#KR1351, KAPA BIOSYSTEMS). The resulting library was sequenced paired-end on an Illumina Nextseq 500 platform. Reads were aligned to the hg38 human genome using the seq2science pipeline (van der Sande et al., 2023), with STAR used as aligner. Reads that mapped equally well to multiple locations were discarded. Count normalization was performed using rlog normalization via DESeq2 in R (Love et al., 2014). Differential genes were considered with adjusted p-value < 0.05 and a fold change > 1.5. For Gene Set Enrichment Analysis, fold changes per gene were determined by calculating the ratio between control and knockdown animals for each biological replicate separately. The average fold change was then used as input for the R package fgsea (Korotkevich *et al*, 2019. *Preprint*).

### Cathepsin labeling

Cathepsin labeling was performed as described before(Edgington-Mitchell et al., 2017). In short, WT or KO gal9 moDCs or isolated WT or KO gal9 CD11c+ splenocytes were harvested and resuspended at 10^6^ cells/mL. Aliquots of 10 µL (0.1e6 cells) were treated with 1 µL 5 µM BMV109 cathepsin probe (final concentration 0.5 µM) and incubated for 1 h at 37 °C. Cells were centrifuged (10,000 rcf, 1 min., RT.), supernatant removed, and the cells were resuspended in 9 µL hypotonic lysis buffer (50 mM PIPES [pH 7.4], 10 mM KCl, 5 mM MgCl_2_, 2 mM EDTA, 4 mM DTT, and 1% NP-40) and incubated 15 min. on ice. Afterwards, lysates were centrifuged (21,130 rcf, 15 min., 4 °C) and supernatant was transferred to fresh tubes. Samples were diluted with 4x Laemmli sample buffer (40% glycerol, Tris/HCl (0.2 M, pH 6.8), 8% SDS, 10% fresh β-mercapto-ethanol, and 0.04% bromophenol blue), denatured at 95 °C for 10 min., and analyzed using 15% SDS-PAGE analysis. Probe signal was visualized using an Amersham Typhoon 5, gel and blot imaging system (Cytiva), followed by equal protein loading confirmation using Trypan Blue staining. Gel analysis was performed using Fiji ImageJ.

### Proteasome labeling

Proteasome labelling was performed as described before (Verdoes et al., 2006). In short, 100,000 moDCs or isolated CD11c^+^ splenocytes were spun down (10,000 rcf, 1 min., RT) and lysed in 9 µL hypotonic lysis buffer for 15 min. on ice. Afterwards, lysates were centrifuged (21,130 rcf, 15 min., 4 °C) and supernatant was transferred to fresh tubes. The lysates were treated with 1 µL 10 µM MV151 (final concentration 1 µM) and incubated for 1 h at 37 °C. Samples were diluted with 4x Laemmli sample buffer, denatured at 95 °C for 10 min., and analyzed using 15% SDS-PAGE analysis. Probe signal was visualized using an Amersham Typhoon 5, gel and blot imaging system (Cytiva), followed by equal protein loading confirmation using Trypan Blue staining. Gel analysis was performed using Fiji ImageJ.

### DQ-OVA endocytosis assay and OVA-presentation assessment

DQ^TM^-OVA (D12053, ThermoFisher) was used as a substrate for assessing DC proteolytic activity. To ensure both WT and galectin-9-depleted moDCs were exposed to the same concentration of substrate, one of the two was stained with Cell Trace Violet^TM^ (1:2000, C34571, ThermoFisher) and mixed in the same well at a 1:1 (WT:KD gal9) ratio of 1x10^5 cells/well. Then, moDCs were incubated with 200 nM Bafilomycin A1 (Sigma-Aldrich) and 0, 5, 10 or 20 μg/mL DQ-OVA for 1.5 h. After incubation, cells were washed extensively and incubated with 1 ug/mL of propidium iodide (in PBS) to differentiate between live and dead cells. Cells were measured using the MACSQuant Analyzer 10 Flow Cytometer (Miltenyi Biotec).

For murine DCs, the antigen processing capacity was determined by incubating the DCs for 24 h with 1 μg/ml LPS and 0, 2.5 or 5 mg/mL OVA (or 400 nM OVA_257-264_ peptide positive control). On the next day, the cells were blocked with 5% mouse serum, incubated with 1 ug/mL of propidium iodide and stained with CD11c (at 1:100 Biolegend; #117311, clone N418, af488-labelled) and H-2Kb bound to SIINFEKL antibodies (at 1:50 Biolegend; #141606, clone 25-D1.16, APC-labelled). Stained cells were analyzed by flow cytometry using a FACSVerse.

### RMA-Muc1 tumor rejection model

*WT, lgals9-/-* and *cd11c^cre^lgals^fl/fl^*female animals were injected subcutaneously with 5×10^6^ RMA-Muc1 cells (as previously described (Dunlock et al., 2022; Gartlan et al., 2013)). Tumor growth was monitored by measuring three dimensions—height (h), width (w), and length (l)—of each tumor using calipers every 2 to 3 days. Tumor volume was then calculated using the formula: π/6 * w*l*h (mm^3^).

### Statistical analysis

Pathway enrichment analysis was conducted using the gProfiler tool (version *e111_eg58_p18_f463989d*). The curated protein list was uploaded to gProfiler, where gene identifiers were matched to their corresponding pathways. The analysis focused on biological pathways (dataset: GO terms or Reactome), molecular functions (dataset: GO terms) and cellular components (dataset: GO terms) relevant to the proteins of interest. The analysis utilized default settings, including statistical significance thresholds set at a p-value < 0.05. Enrichment results were interpreted using Benjamini-Hochberg correction for multiple testing. Pathways were considered significantly enriched if they met the specified thresholds. The results were visualized using R to illustrate the relationships and significance of the identified pathways.

All data was processed using Excel 2019 (Microsoft) and plotted using GraphPad Prism 8 or 10 (Version 8.0.2. or 10.0.0 GraphPad Software). Unless otherwise stated, all data is expressed as mean +/- SEM. Significance among the different conditions was assessed by one-way ANOVA with Tukey multiple comparison corrections, two-way ANOVA with Šídák’s multiple comparisons test, or an unpaired or paired t-test. Statistical significance threshold was defined as *p < 0.05; **p < 0.01; ∗∗∗p < 0.001. All experiments were performed in duplicate to ensure reproducibility.

## Results

### Downregulation of galectin-9 affects transcriptional programs associated with antigen presentation

We examined whether gal9 loss resulted in transcriptional changes in DCs using mature non-targeting siRNA or *Lgals9* siRNA-transfected monocyte-derived DCs (referred to as WT and KD gal9 DCs, respectively) (Fig. 1A). Gal9 protein surface expression was almost completely inhibited by this approach, while DC maturation (analyzed by HLA-DR, HLA-ABC, CD80, CD70 and PD-L1 protein expression) was not affected by gal9 depletion (Fig. S1A - S1B). We then performed RNA-seq analysis of WT and KD gal9 DCs from three independent donors and observed overall transcriptional changes as a result of gal9 downregulation, with gal9 being among the most downregulated genes (Fig. 1B). To study the biological effect of the knockdown, we performed gene set enrichment analysis. To account for inter-donor variation in gene expression as readily observed for the identified differential genes (Fig. S2A), we used the average fold change between the replicates as input (Fig. 1C). This revealed that gal9 knockdown results in increased expression of genes involved in antigen processing via MHC-II, and reduction of gene expression programs involved in T cell proliferation and activation (Fig. 1D, Fig. S2B). Together, these data indicate that gal9 in DCs plays a role in the antigen presentation-recognition axis.

**Figure 1.**
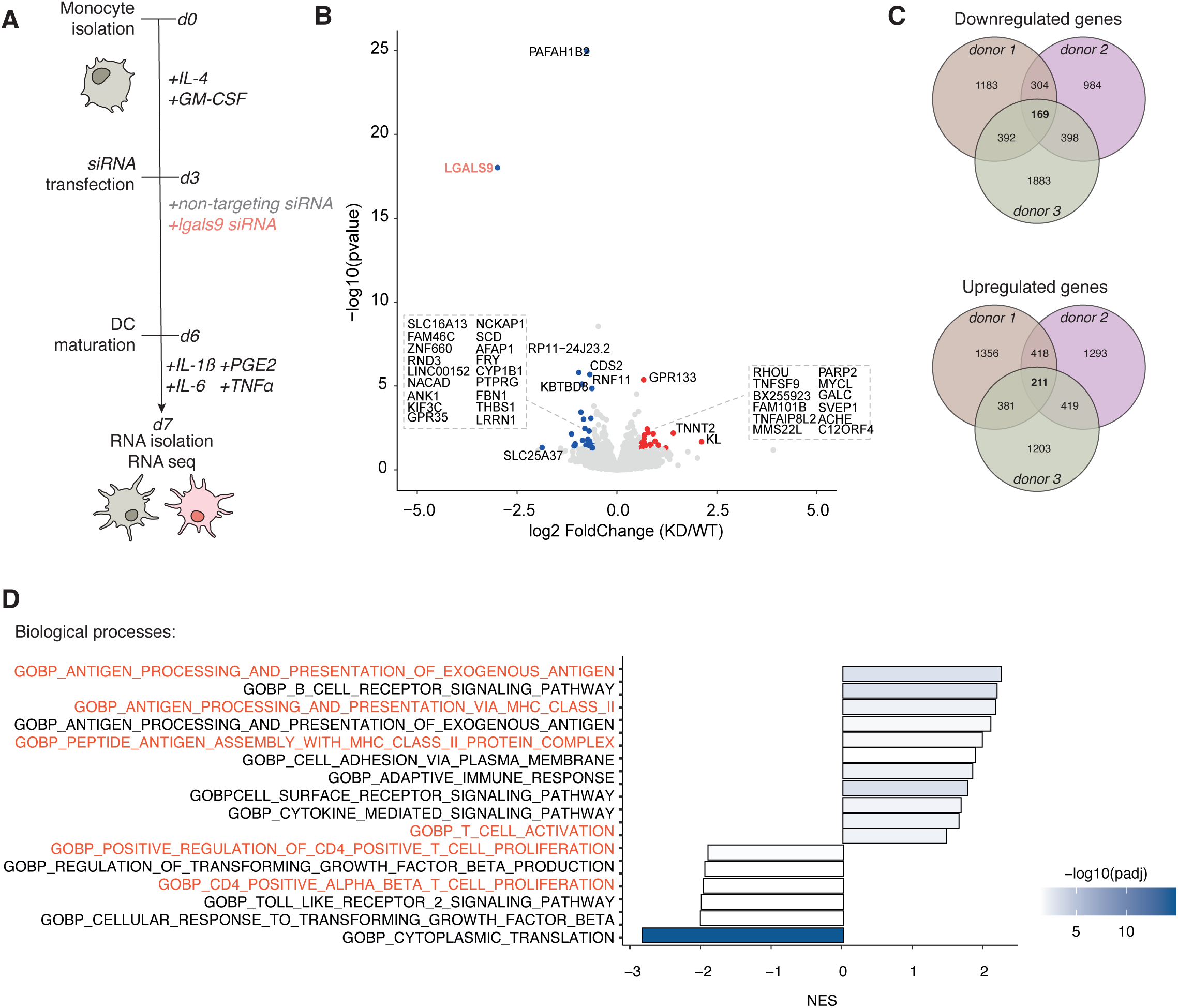
Gal9-depleted moDCs display altered T cell proliferation and MHC-II complex assembly pathways. **(A)** Graphical representation of the experimental timeline for RNA sequencing of WT and gal9 KD moDCs. moDCs were generated by isolating CD14^+^ cells from total PBMCs and treated with IL-4 and GM-CSF for three days prior to being transfected with a non-targeting or a *lgals9* siRNA. After day 6 of culture moDCs were matured with a mixture of pro-inflammatory cytokines (IL-6, IL-1β, TNFɑ and PGE2) for 24 h after which total RNA was isolated for sequencing analysis. **(B)** Volcano plot of differentially expressed genes comparing KD gal9 to WT conditions. Red dots denote significantly upregulated genes, and blue dots significantly downregulated genes. Differentially expressed gene names are indicated next to the dot or in a squared gray dotted box. Fold change represents KD gal9 vs. WT moDC expression levels. **(C)** Venn diagrams showing overlap in gene expression changes among the tree donors analyzed. The top diagram represents the number of genes commonly upregulated across all donors, while the bottom diagram displays downregulated genes. **(D)** Gene set enrichment analysis using gene ontology (GO) terms, from the Biological Process dataset performed on gProfiler. Color intensity corresponds to the -log adjusted p-value, with darker colors indicating greater significance in pathway enrichment. MHC-II and T cell activation and proliferation-related pathways are highlighted in red.

### Galectin-9 is required for CD4^+^ T cell activation

We then investigated the functional consequences of gal9 depletion in DC-mediated T cell activation using a mixed lymphocyte reaction (MLR) in which WT or KD gal9 moDCs were co-cultured with allogeneic T cells. Depletion of gal9 in DCs diminished T cell activation as seen by the lower expression of the activation marker CD25 at the surface of T cells upon co-culture with KD gal9 DCs (Fig. 2A and 2B). Concomitantly, these T cells also exhibited lower IFN-γ secretion (Fig. 2C) and proliferation capacity (Fig. 2D and 2E) when compared to T cells activated by WT DCs.

**Figure 2.**
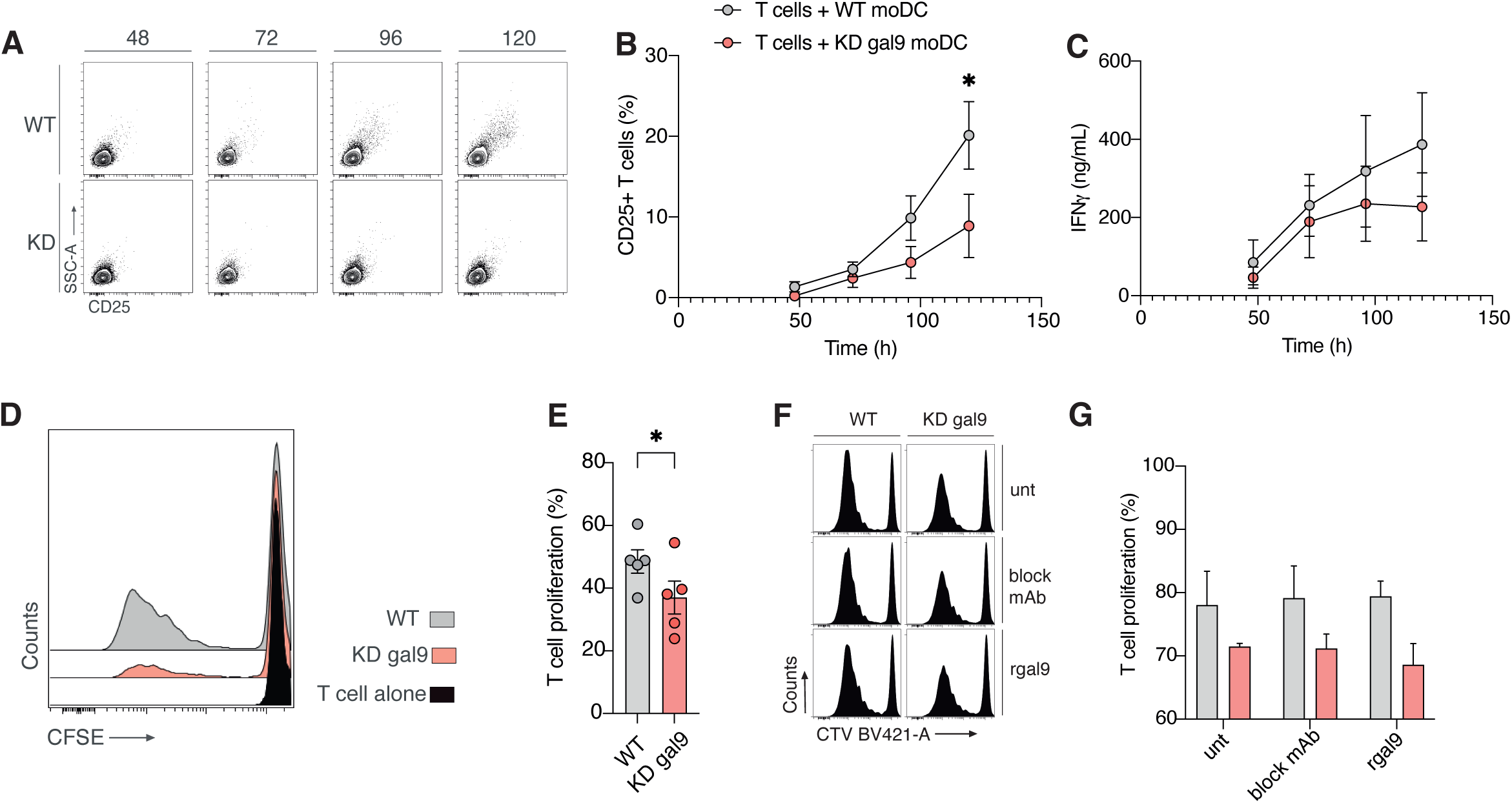
Intracellular gal9 regulates DC-mediated allogeneic T cell effector function. **(A)** Representative flow cytometry dot plot depicting CD25 (activation marker) surface staining in T cells cultured with either human WT or KD gal9 DCs for 48, 72, 96 and 120 hours. **(B)** Quantification of data shown in (A) from four independent donors. Graph depicts dot plot CD25 expression in T cells cultured with human WT or KD gal9 DCs for 48, 72, 96 and 120 hours. **(C)** IFNγ (ng/mL) secreted by T cells cultured with human WT or KD gal9 DCs for 48, 72, 96 and 120 hours. **(D)** Flow cytometry histogram showing proliferating allogeneic CFSE-labeled T cells cultured alone (black) or together with either WT or KD gal9 DCs. **(E)** Quantification (percentage of proliferating T cells) of data shown in (D) for 5 independent donors. **(F)** Representative proliferation histogram of T cells cultured with WT or gal9 KD DCs untreated or treated with either anti-gal9 blocking antibody or with recombinant gal9 for 1 hour. **(G)** Percentage of allogeneic CTV-labeled proliferating T cells of data shown in (F) from two independent donors. Gray = T cells co-cultured with WT DCs. Red = T cells co-cultured with KD gal9 DCs. Data represent mean ± SEM of 3-5 independent donors. Two-way ANOVA with Šídák’s multiple comparisons test or paired t-test was conducted between conditions. *p < 0.05.

Since gal9 is located both in the cytosol and extracellularly (membrane-bound) in DCs (Querol Cano et al., 2019), we investigated the involvement of both subcellular fractions in driving T cell activation by treating KD gal9 DCs with exogenous recombinant gal9 (rGal9) prior to subjecting them to MLR assays. In addition, WT DCs were treated with a specific gal9 blocking antibody (Dai et al., 2005). Analysis of gal9 expression revealed that exogenous protein (rGal9) restored surface-bound levels of gal9 (Fig. S3A). However, treatment with rGal9 did not rescue the impairment in T cell activation and concomitantly, blocking of extracellular gal9 in WT DCs did not detrimentally alter T cell proliferation indicating that the intracellular fraction of gal9 is responsible for the diminished T cell activation (Fig. 2F and 2G).

Next, we validated gal9 involvement in T cell activation using autologous antigen-driven models as they pose a more physiological setup (Fig. 3A). First, we studied CD4^+^ T cell activation using tetanus toxoid (TT) as antigen. WT DCs significantly induced TT-specific T cell proliferation whereas this was significantly inhibited by T cell co-culture with KD gal9 DCs (Fig. 3B). Interestingly, non-specific CD4^+^ T cell proliferation was also diminished in the presence of KD gal9 DCs suggesting gal9-induced effects on T cell proliferation are not antigen specific (Fig. 3B). In contrast, autologous proliferation assays performed using gp100 (280-288) peptide, a CD8^+^ T cell specific antigen (Fig. 3A), revealed no effect of gal9 depletion in T cell activation. No differences in T cell proliferation were observed when T cells transfected with the specific gp100_TCR (Fig. S3B) were co-cultured with either WT or KD gal9 DCs and in the presence of the relevant gp100 peptide (Fig. 3C). In agreement, the expression of perforin and granzyme B in gp100_TCR^+^ T cells as well as their TNFα and IL-2 production markedly increased upon activation with gp100-pulsed DCs regardless of their gal9 expression (Fig. S3C). Similarly, the expression of the activation markers CD25, CD69 and CD137 increased in gp100_TCR-transfected T cells upon co-culture with gp100-presenting DCs but was comparable between WT and KD gal9 DCs (Fig. S3D). T cell co-cultures with untreated DCs or with DCs pulsed with an irrelevant peptide induced no T cell proliferation or activation confirming the specificity and robustness of our data (Fig. 3C, S3C and S3D). Thus, these data indicate that gal9 on human DCs is required for CD4+ T cell proliferation.

**Figure 3.**
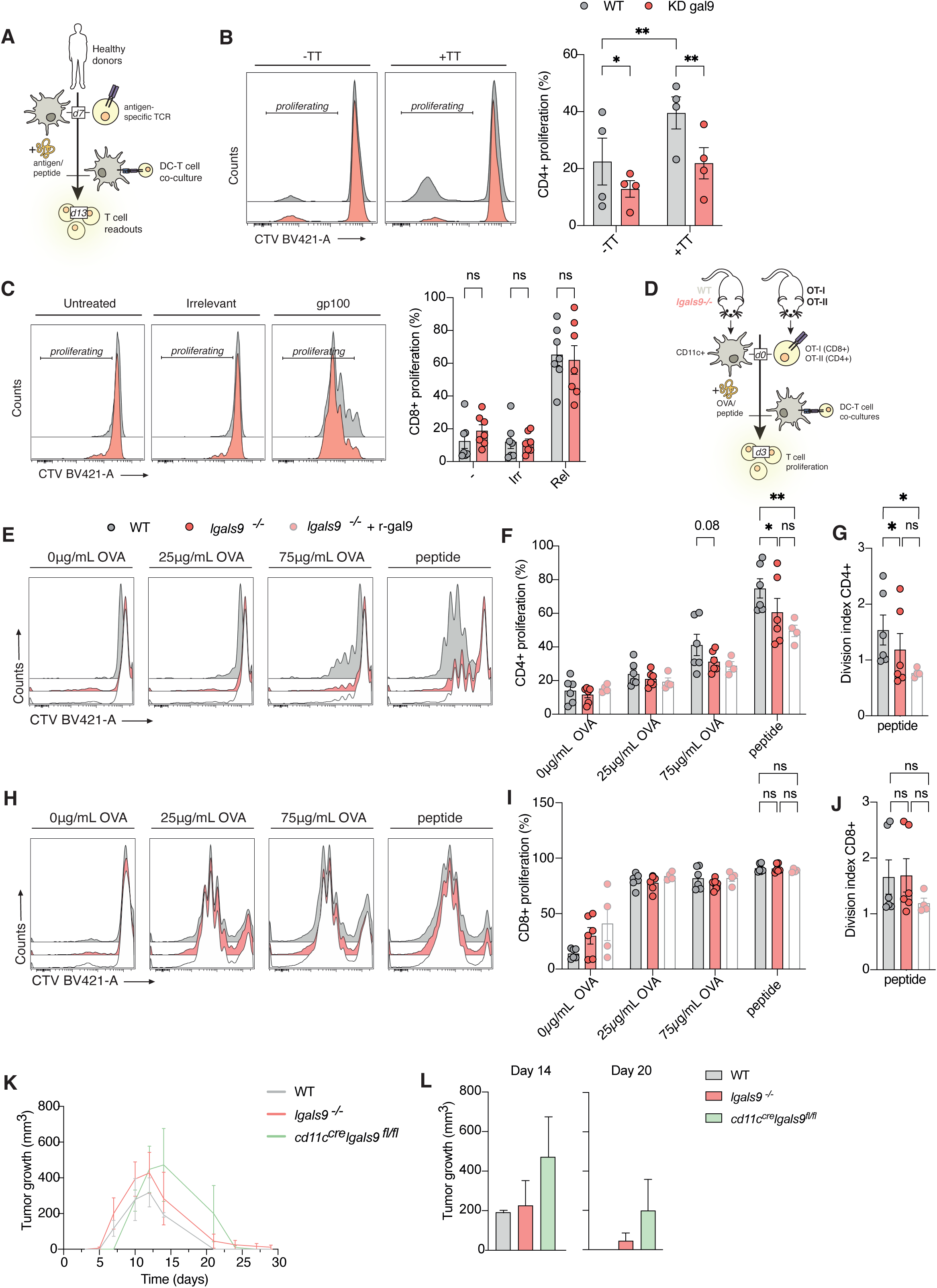
Gal9-deficient-DCs are impaired in their ability to induce CD4^+^ T cell proliferation. **(A)** Graphical representation of the human-based experimental pipeline. Both monocytes and T cells were isolated from PBMCs obtained from healthy donors. Monocytes were differentiated into moDCs and primed with the (irr)relevant antigen or peptide. In parallel, autologous T cells were transfected with an mRNA encoding for gp100-TCR or obtained from vaccinated donors with Tetanus toxoid (TT)-TCR specific clones. Cells were co-cultured for seven days after which T cell proliferation was measured. **(B)** (*left*) Human WT (grey) or KD gal9 (red) DCs treated with tetanus toxoid antigen (+TT) or PBS (-TT) as negative control were incubated with autologous CD4^+^ T cells. Representative flow cytometry histogram plots depict proliferation peaks of CTV-labeled CD4^+^ T cells co-cultured with DCs in the presence or absence of TT treatment. (*right*) Quantification of CD4^+^ T cell proliferation shown in the left panel (B). Each dot represents an independent donor (n=4). **(C)** (*left*) Human WT or KD gal9 DCs were treated with either PBS (−), irrelevant peptide (NY-ESO, Irr), and relevant peptide (gp100, Rel) prior to being co-cultured with autologous CD8^+^ T cells. Plots depict flow cytometry histogram of CTV-labeled CD8^+^ T cells. (*right*) Quantification of CD8^+^ T cell proliferation from data shown in left panel (C). Each dot represents an independent donor (n=7). **(D)** Graphical representation of the murine-based experimental pipeline. WT and *lgals9*^-/-^ CD11c^+^ cells were obtained from their respective mice strains. T cells were obtained from OT-I or OT-II transgenic mice. CD11c^+^ cells were treated with LPS and OVA or its peptides. Cells were co-cultured for three days after which T cell proliferation was measured. **(E)** WT (grey), gal9 knockout (*lgals9* ^-/-^, red), or *lgals9* ^-/-^ DCs treated with exogenous recombinant murine gal9 (*lgals9*^-/-^ + r-gal9, white) were pulsed with 0, 25, or 75 μg/mL OVA protein or with a MHC-II-restricted peptide prior to being co-cultured ex vivo with OT-II T cells. Data depicts representative flow cytometry histogram of CTV-labeled OT-II T cells. **(F)** Quantification of OT-II CD4^+^ T cell proliferation from data shown in (E) (n= 4-6 mice/genotype from three individual experiments). **(G)** Division index of the parental OT-II CD4^+^ T cells treated as in (E), representing the number of divisions in subsequent generations upon incubation with DCs. **(H)** Representative histogram plots of OT-I CD8^+^ T cell proliferation in murine DC cultures. WT, gal9 KO, or gal9 KO +r-gal9 DCs were treated with 0, 25, or 75 μg/mL OVA protein or with an MHC-I-restricted peptide prior to being co-cultured with OT-I T cells. **(I)** Quantification of OT-I CD8^+^ T cell proliferation shown in (H) (n= 4-6 mice/genotype from three individual experiments). **(J)** Division index of the parental OT-I CD8^+^ T cells treated as in (H), representing the number of divisions in subsequent generations upon incubation with DCs. **(K)** RMA-Muc1 tumor growth (in mm^3^) in WT (gray)*, lgals9^-/-^* (red) and *cd11c^cre^lgals9^fl/fl^* (green) mice over time. **(L)** Tumor volume for WT*, lgals9^-/-^* and *cd11c^cre^lgals9^fl/fl^* mice on days 14 and 20 post tumor cell injection. n = 3-4 mice/genotype (mean ± SEM, Two-way ANOVA with Tukey’s multiple comparisons test). All bar graphs show mean ± SEM. Each dot represents one independent donor (human data) or mouse. Two-way ANOVA with Šídák’s multiple comparisons test was conducted between conditions. *p < 0.05, **p < 0.005.

Next, we investigated whether gal9 function in T cell activation was conserved across species by using ovalbumin (OVA) in T cell proliferation assays on splenic DCs (CD11c^+^) isolated from WT and gal9 ^-/-^ (KO) mice. CD8^+^ and CD4^+^ T cells were harvested from TCR transgenic OT-I and OT-II mice respectively and stimulated *in vitro* with WT or gal9 KO DCs primed with either increasing concentrations of OVA or peptide (Fig. 3D). CD4^+^ T cell proliferation was reduced in CD4^+^ T cells activated by gal9 KO DCs compared to WT DCs (Fig. 3E – 3G), whereas CD8^+^ T cell proliferation was not affected by gal9 depletion (Fig. 3H – 3J) confirming that gal9 selectively impairs CD4^+^ T cell activation. The role of gal9 in T cell proliferation was confirmed by quantifying the division index (the number of divisions each T cell underwent), which was significantly lower in CD4^+^ T cells co-cultured with gal9 KO DCs compared to their WT counterparts. In contrast CD8^+^ proliferation was not affected by gal9-deficiency (Fig. 3G and 3J) in line with human DC data. In addition, treatment of gal9 KO DCs with rGal9 did not rescue the impairment in total CD4^+^ T cell proliferation nor in the average cell division index (Fig. 3E – 3G), whereas treatment with rGal9 had no effect on CD8^+^ T cell proliferation (Fig. 3H – 3J). Together, these data demonstrate that intracellular gal9 in DCs is responsible for CD4+ T cell activation and proliferation.

### DCs from gal9 KO mice are impaired in tumor rejection capacity *in vivo*

Since gal9 is also expressed in other immune cells, we generated gal9 conditional knockout mice to specifically dissect the effects of gal9 in DCs *in vivo*. *Lgals9*-floxed (*lgals9^fl/fl^*) mice were crossed with *cd11c^cre^* transgenic mice to generate DC-gal9 conditional KO mice (*cd11c^cre^lgals9^fl/fl^*). Characterisation of the myeloid compartment of WT, *gal9^-/-^* or *cd11c^cre^lgals9^fl/fl^* mice demonstrated no impairment in DC development upon gal9 loss (Fig. S4 and Fig. S5A-5J). Similar numbers of conventional DC type 1 (cDC1), plasmacytoid DC or conventional DC type 2 (cDC2) were observed in lymph organs of WT, *gal9^-/-^* or *cd11c^cre^lgals9^fl/fl^*mice (Fig. S5A-5E), and migratory cDC1 and cDC2 subsets were also found in comparable numbers (Fig. S5B and S5D). In addition, no differences in the percentage of macrophages, eosinophils, neutrophils or monocytes were observed between WT, *gal9^-/-^* or *cd11c^cre^lgals9^fl/fl^* mice (Fig. S5F-S5J), implying that gal9 is not required for myeloid cell development and differentiation. Next, we investigated the T cell-dependent tumor rejection model RMA-Muc1 (Dunlock et al., 2022; Plunkett et al., 2004) to determine whether gal9 expression in DCs was relevant to T cell function *in vivo*. Loss of gal9 in DCs delayed anti-tumor responses as tumor burden in *cd11c^cre^lgals9^fl/fl^*mice was larger as compared to WT and *gal9*^-/-^ mice after day 12 and until day 22 post tumor inoculation (Fig. 3K and 3L). These data indicate that gal9 expression in DCs controls T cell anti-tumor activity *in vivo*.

### Immune synapse formation is dependent on galectin-9

To elucidate the mechanism behind impaired T cell activation upon gal9 loss, we first examined whether gal9 modulates antigen processing and presentation. First, human WT or KD gal9 DCs were treated with increasing concentrations of self-quenching OVA (DQ-ova) (Pascottini et al., 2019), which emits fluorescence upon proteolytic degradation. The extent of antigen processing, in the form of cleaved DQ-ova, was evaluated by flow cytometry and revealed gal9 is not involved in antigen processing as no differences in fluorescence signal were observed between WT or KD gal9 DCs when cells were pulsed with increasing DQ-ova concentrations (Fig. 4A and 4B). Pre-treatment with Bafilomycin A, a lysosome inhibitor that blocks endosomal acidification, resulted in a complete abrogation of antigen processing in both WT and KD gal9 DCs, confirming the pH-dependency of degradation (Fig. 4A and 4B). To further corroborate depleting gal9 does not compromise the proteolytic capacity of DCs, we examined cysteine cathepsin activity in WT and gal9-depleted murine and human DCs using the pan-reactive activity-based probe BMV109. As shown, incubation with the probe resulted in equal labeling intensities for all cysteine cathepsins detected in both WT and gal9-depleted cells (Fig. S6A and S6C). Similarly, treatment with MV151, an activity-based probe targeting the immuno- and constitutive proteasome also revealed equal activity of labelled catalytic β and βi subunits (Fig. S6B and S6D) in both human and murine gal9-depleted DCs compared to their WT counterparts. Next, we pulsed murine WT or gal9 KO DCs with increasing concentrations of OVA and determined MHC-peptide occupancy by flow cytometry. No differences were observed in antigen binding to MHC in WT or gal9 KO DCs showing that gal9 is not required for antigen presentation itself (Fig. S6E and S6F). Altogether, this data demonstrates that gal9 is not involved in antigen processing or presentation.

**Figure 4.**
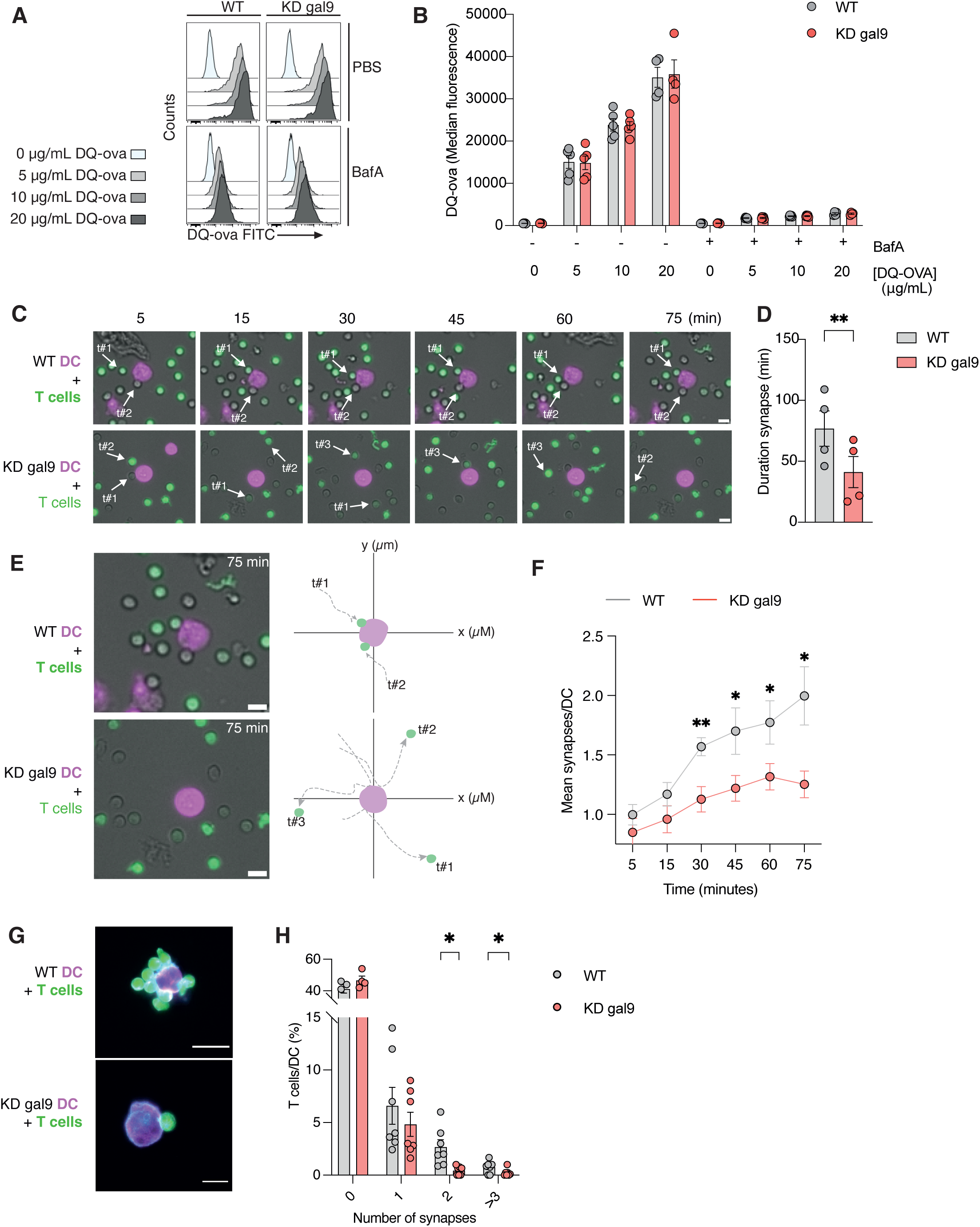
Galectin-9 deficient DC fail to establish stable interactions with T cells. **(A)** Flow cytometry histograms depicting the fluorescence intensity of the DQ-OVA probe (0, 5, 10, 20 μg/mL) in wild type (WT, grey) or gal9 depleted DCs (KD gal9, red) treated with PBS or Bafilomycin A (BafA) as inhibitor for lysosome acidification. **(B)** Quantification of data shown in (A) from five independent donors. Gray: WT DCs. Red: KD gal9 DCs **(C)** Time-lapse images of WT or KD gal9 DCs co-cultured with T cells for 5, 15, 30, 45, 60, and 75 minutes showing DC-T cell contacts. DCs are shown in magenta and T cells in green. White arrows illustrate the tracking of specific T cells towards a DC. **(D)** Quantification of the time (minutes) during which T cells establish a synaptic contact with a DC from the time lapses depicted in (C). Analysis done from the mean of 10-15 cells from 4 independent donors (*left*) or 36 representative synapses (*right*). **(E)** Representative images captured at 75 minutes of DC-T cell co-cultures (*left*) together with a graphic illustration of the migration tracks from T cells over time (*right*). DCs are depicted in magenta and T cells in green. **(F)** Quantification of the amount of DC-T cell contacts over time from the time lapses shown in (C). Analysis done for nine independent donors. **(G)** Allogeneic WT and KD gal9 DCs (magenta) were co-cultured with T cells (green) for 2 hours in low-adhesion plates. Following incubation, cells were fixed and transferred to an imaging plate for visualization. **(H)** Quantification of the number of WT or KD gal9 DC establishing contact with allogeneic T cells (0, 1, 2 or >3 T cells/DC) of data shown in (G) for 4-7 independent donors. In D, F and H; Gray: T cells co-cultured with WT DCs. Red: T cells co-cultured with KD gal9 DCs. Scale bars = 10 µm. All graphs depict mean values ± SEM. Two-way ANOVA with Šídák’s multiple comparisons test was conducted between conditions (for B and H) unpaired T-test analysis (for D and F). *p < 0.05, **p < 0.005.

Next, we determined the ability of WT and KD gal9 DCs to interact with T cells using an allogeneic DC-T cell synapse model with superantigen SEB (Fig. S7A). Gal9 depletion decreased the ability of DCs to establish stable cell-cell contacts with T cells (Fig. 4C). Interactions between WT DCs and T cells were long-lasting with an average duration of approximately 1 h whereas KD gal9 DC – T cell interactions were transient, lasting only about 20 min (Fig 4D). Individual T cell tracks depict the enhanced motility of T cells in the vicinity of KD gal9 DCs compared to their suppressed locomotion, subsequent of T cell receptor engagement (Dustin, 2014), when in contact with WT DCs (Fig. 4E). In addition, WT DCs established simultaneous synaptic interactions with multiple T cells over time, whereas gal9 depletion rendered DCs unable to do so (Fig. 4F). This was confirmed using an MLR model in which KD gal9 DCs were significantly impaired in their ability to establish contacts with more than one T cell at a time (Fig. 4G and 4H). In contrast, WT DCs displayed a significantly enhanced capacity to form so-called rosette structures with multiple T cells, necessary for proper T cell activation (Bousso and Robey, 2003). Thus, these data show that gal9 in DCs is required for immunological synapse formation.

### Galectin-9 recruits HLA-DR to the immune synapse between dendritic cells and T cells

To mechanistically resolve how gal9 is involved immune synapse formation we examined whether gal9 was located in the contact zone between DCs and T cells using airyscan confocal microscopy. Gal9 was distributed uniformly in naïve DCs but specifically recruited to the immune synapse upon T cell interaction (Fig. 5A). Gal9 enrichment quantification demonstrated increased localisation of gal9 at the DC side of the IS, indicating a putative role for gal9 in IS assembly (Fig. 5B).

**Figure 5.**
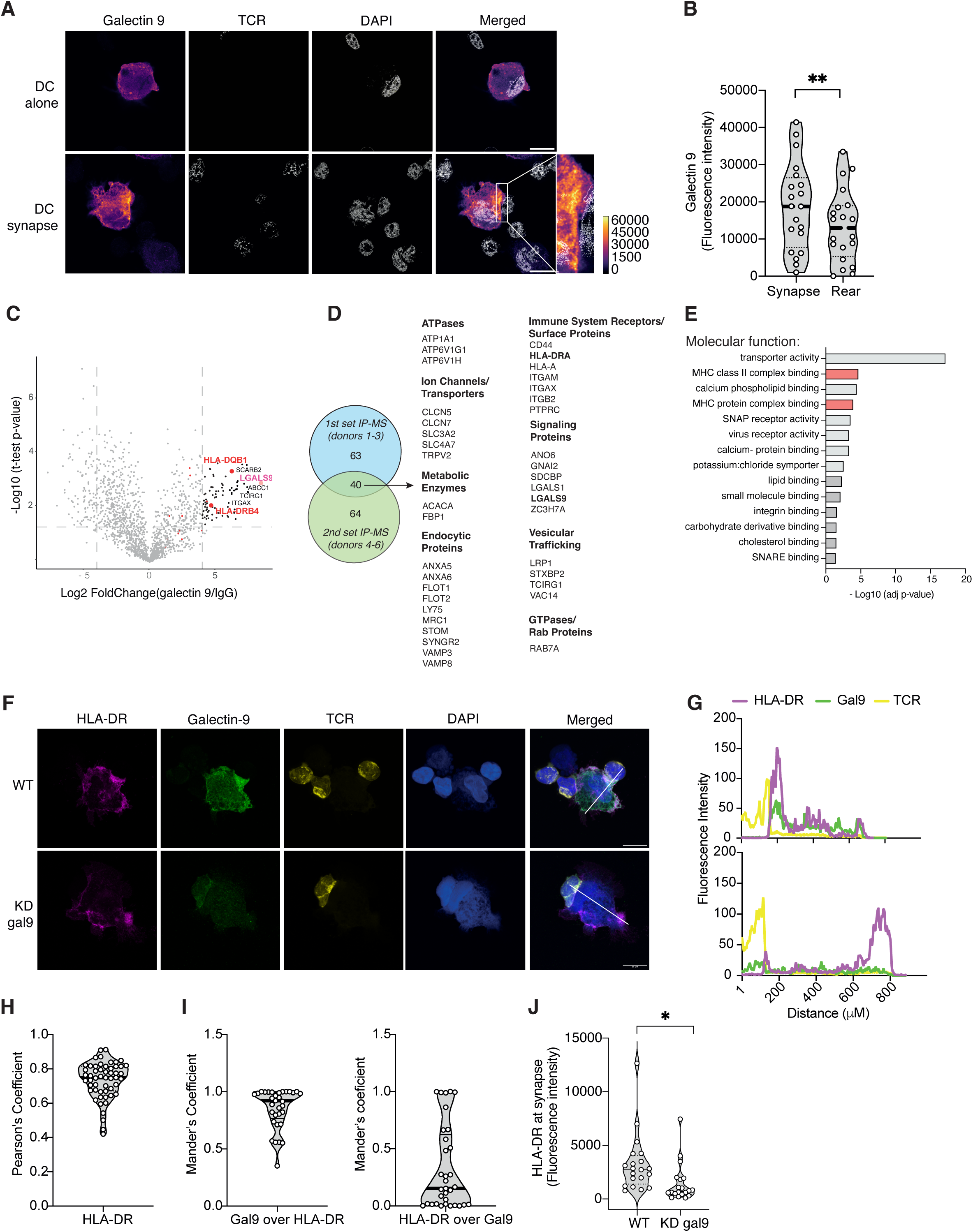
Galectin-9 recruits and colocalizes with HLA-DR at the DC-T cell immune synapse. **(A)** Representative microscopy images of DCs alone (top) or in co-culture with T cells (bottom). Cells were stained against gal9, TCR and DAPI (nuclear marker). A merged composite image is also shown. Fluorescence intensity isdepicted as colorized pixels, with grayscale values representing signal intensity. The zoomed-in image (2x) highlights the detailed localization of gal9 at the DC-T cell interface. **(B)** Quantification of fluorescence intensity values of gal9 at the immune synapse compared with the rear of WT DCs co-cultured with T cells. Graph depicts mean ± SEM for three independent donors (6-10 cells per donor). Unpaired T-test analysis was performed, **p < 0.005. **(C)** Volcano plot depicting gal9 interacting proteins. Differentially enriched proteins binding to gal9 (right) or isotype (left) are shown. MHC-II (HLA-DR and HLA-DQ) molecules are highlighted in red. Dotted gray lines depict the cut-off values; the fold change (x axis) of 2 and p-value (y axis) of 0.05. Black dots represent the proteins that had fold change >2 and p-value >0.05. **(D)** Venn diagram showing the overlap of gal9 binding proteins identified from two sets of mass spectrometry-immunoprecipitation experiments, each performed with three independent donors. A total of 40 common hits independently enriched at least 1.5 fold between gal9 and isotype pull downs are shared between the two experiments. The list of gal9 binding proteins (gene names) is categorized by their respective biological functions. **(E)** Pathways significantly enriched related to gal9 interacting proteins (from the molecular function database). Pathways related to MHC-II function are highlighted in red. Statistical significance is represented by the adjusted p-value (-log10) (x axis). **(F)** Representative microscopy images of WT or gal9 KD DCs incubated for 2h with T cells before being fixed and stained for HLA-DR, gal9, TCR and DAPI (nuclear marker). The white line in the merged composite image indicates the arbitrary line that was used to depict the line scans in (G). **(G)** Representative line scans of cells shown in (F) depicting HLA-DR (magenta), TCR (yellow) and gal9 (in green) intensity across the line (starting at the synapse and finishing at the rear of the cell). **(H)** Pearsońs and Mandeŕs **(I)** correlation coefficient for HLA-DR and gal9 was calculated from images shown in **(F)**. 53 WT DCs were quantified from 3 independent donors. **(J)** Quantification of HLA-DR fluorescence intensity at the synaptic site in WT and KD gal9 DC co-cultured with T cells. Scale bars = 10 µm. Unpaired student’s t-test was conducted between WT and KD gal9 conditions. Data represent mean ± SEM of three independent donors. 6-10 cells were analyzed for each condition per donor. *p < 0.05

Next, we performed immunoprecipitation coupled with mass spectrometry (IP-MS) to identify gal9 binding partners in DCs. Gal9 was significantly enriched in the gal9-IP compared to isotype control, confirming the quality of the pull-down (Fig. 5C). To account for donor variability, we performed two independent IP-MS with DCs isolated from six independent donors (Fig. 5D). Forty binding partners were found to be enriched at least 1.5-fold in gal9 pull down compared to isotype control in both IP-MS experiments, including previously reported interactions (e.g. CD44, Vamp3) (Fig. 5D) (Santalla Mendez et al., 2023). Analysis of gal9-binding proteins identified human leukocyte antigen-class II (HLA-DR) to interact specifically with gal9 (Fig. 5C and 5D). Gene Ontology (GO) analysis of the gal9 binding partners showed a functional enrichment in MHC-class II protein complex binding (Fig. 5E and Fig. S7B). HLA-DR (MHC-II) is responsible for presenting antigens to CD4^+^ T cells and thus we further examined whether gal9 interaction with HLA-DR underlies the impaired IS formation in KD gal9 DCs. We observed co-localisation of HLA-DR and gal9 at the DC side of the immune synapse (visualised by the localisation of the TCR) using airyscan confocal microscopy (Fig. 5F and 5G), which was confirmed by quantifying Pearson’s (Fig. 5H) and Mander’s (Fig. 5I) correlation coefficients. This colocalization was only observed after cell permeabilization (data not shown), which supports our previous finding that intracellular gal9 is responsible for driving IS formation. Notably, depletion of gal9 in DCs impaired HLA-DR recruitment to the interaction zone with T cells (Fig. 5F and 5G). Quantification of HLA-DR expression by fluorescence microscopy demonstrated that HLA-DR recruitment to the immune synapse was significantly decreased in KD gal9 DCs compared to WT cells confirming a functional relationship between HLA-DR and gal9 (Fig. 5J).

Overall, these results demonstrate gal9 depletion disrupts HLA-DR recruitment to the IS thus establishing gal9 as essential for synapse formation and subsequent T cell activation and proliferation.

## Discussion

Dendritic cells are crucial for initiating cellular immunity by activating T cells through MHC-mediated antigen presentation, a process dependent on the formation of the IS (Banchereau and Steinman, 1998). While the molecular mechanisms driving the assembly of the IS at T cell side have been well-studied, how the IS components are trafficked and assembled at the plasma membrane of DCs remains elusive.

In this study we identified gal9 is required for IS formation and report a novel functional interaction with MHC-II (HLA-DR). We found that depletion of gal9 in DCs disrupted the localization of HLA-DR at the synapse, hindering stable DC-T cell interactions and impairing CD4^+^ T cell proliferation. *In vivo*, the absence of gal9 in DCs resulted in increased tumor growth, underlining its importance in DC-mediated immune responses. Our findings demonstrate that gal9 enables sustained DC-T cell contacts and effective CD4^+^ T cell proliferation (Fig. 6).

**Figure 6.**
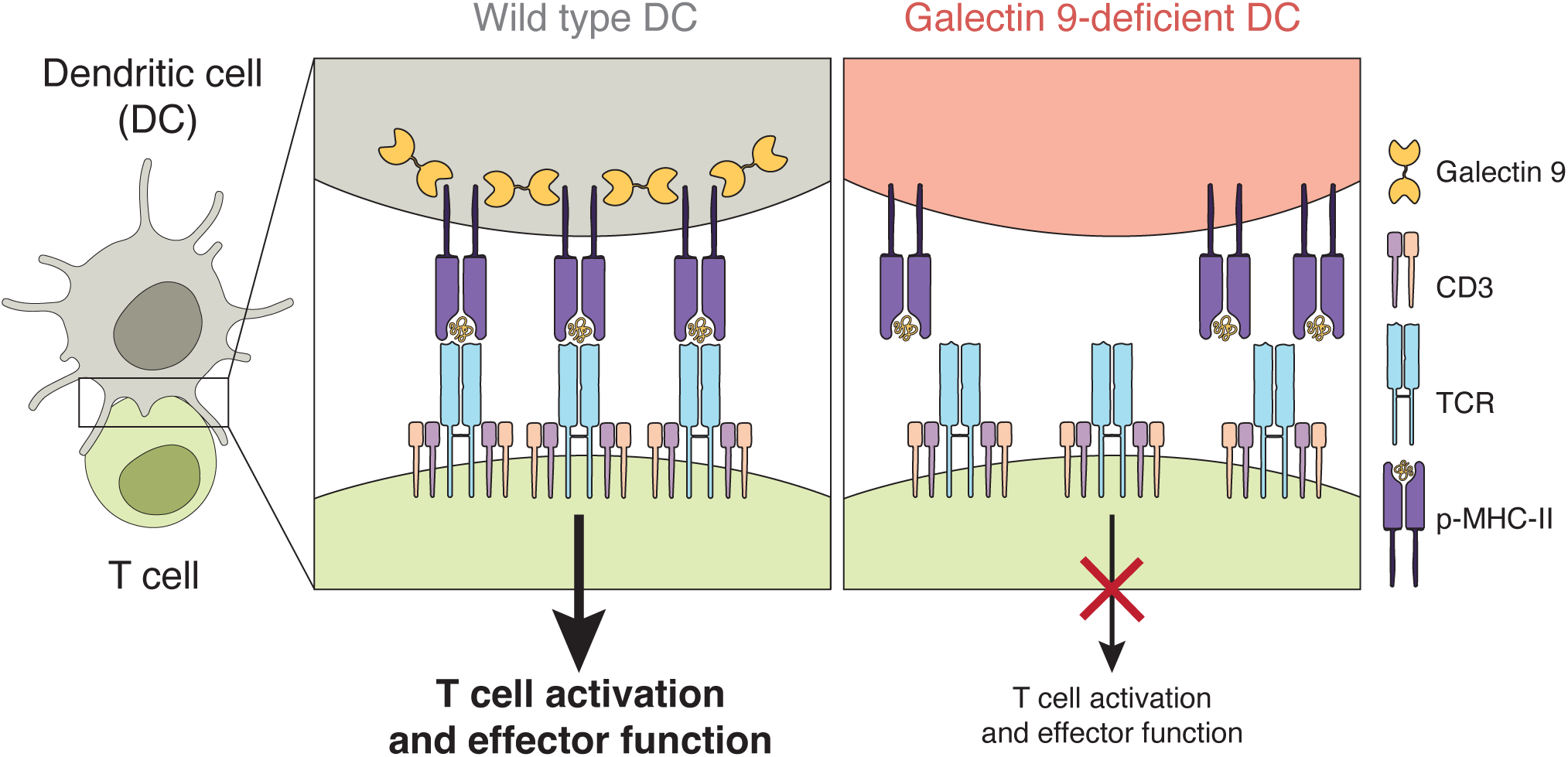
Graphical abstract. Galectin-9 enables immune synapse formation between DCs and T cells. (L*eft*) Galectin-9 binding to HLA-DR drives its recruitment to the immune synapse, enabling efficient DC-T cell engagement resulting in T cell activation and effector function. (*Right*). In the absence of galectin-9, HLA-DR fails to be recruited to the immunological synapse, impairment DC-mediated T cell activation

Gal9 is mostly known as the ligand of the T cell inhibitory receptors TIM-3 and PD-1 through which it induces T cell apoptosis (Yang et al., 2021). Similarly, gal9 suppresses B cell activation by directly binding to the B cell receptor and enhancing interactions with inhibitory co-receptors CD45 and CD22 (Cao et al., 2018; Giovannone et al., 2018). Contrary to its immunosuppressor functions in lymphocytes, we and others have reported a stimulatory function for gal9 in DCs. For instance, addition of recombinant galectin-9 induced the maturation of DCs (Dai et al., 2005) and we have demonstrated gal9 is necessary for optimal phagocytic capacity and cytokine secretion in DCs (Querol Cano et al., 2019; Santalla Mendez et al., 2023). Data presented here highlights a previously unreported role for gal9 in ensuring DC-mediated T cell activation, with consequences for launching cellular adaptive immune responses. Our findings are in line with recent single cell RNA sequencing data showing that high gal9 expression in DCs correlated with proficient antigen presentation and co-stimulatory molecule expression as well as efficient T cell interactions (Meiser et al., 2023). Moreover, DC conditional gal9-KO mice exhibited increased tumor growth compared to both the full gal9 KO and WT counterparts using the T cell-dependent tumor rejection model RMA-Muc1, which underscores the critical role of gal9 in T cell-mediated anti-tumor immunity. The discrepancy between the conditional and the total KO model in rejecting the tumor may be explained by the cell specific anti- and pro-inflammatory functions of gal9, due to the distinct binding partners expressed by different immune cells that can vary based on the differentiation and activation state.

Galectins have been shown to mediate T cell function and T cell glycosylation repertoire changes upon activation, in turn affecting galectin-mediated interactions and receptor dynamics (Demetriou et al., 2001). For instance, galectin-3 (gal3) organizes the N-glycosylated TCR lattice on T cells, inhibiting spontaneous oligomerization, and preventing unintended signaling in naïve cells (Demotte et al., 2008; Petit et al., 2016). Intracellularly, several galectins have also been shown to regulate IS formation in T cells. Illustrating this, gal3 translocates to the cytosolic side of the IS in CD4^+^ T cells upon activation, where it downregulates the surface expression of key TCR components, thereby limiting T cell activation (Chen et al., 2009). Conversely, gal9 is recruited to the intracellular side of the IS in T cells during activation, where it enhances T cell activation by influencing TCR-CD3 complex formation and calcium mobilization (Chen et al., 2020; Lhuillier et al., 2015). While the roles of galectins at the T cell synaptic site have been extensively studied, the molecular mechanisms by which galectins regulate DC functions at the IS remain virtually unknown.

Our data demonstrate an intracellular function for gal9 in driving MHC-II recruitment to the IS. Remarkably, HLA-DR membrane expression and upregulation upon maturation were not altered in response to gal9 depletion, suggesting gal9 is required for the assembly of HLA-DR-membrane microdomains upon DC – T cell interaction. Supporting this hypothesis, we observed no differences in cathepsin expression or activity within endocytic compartments in human or murine gal9-depleted DCs. Given that cathepsins, particularly Cathepsin S, are crucial for the maturation of HLA-DR during the processing of the invariant chain (Bania et al., 2003; Roche and Cresswell, 1991), our findings indicate that gal9 does not influence antigen processing or peptide loading in the endocytic compartment. Concomitantly, our data revealed no differences in antigen processing and presentation between gal9 depleted DCs and their WT counterparts. Thus, the intracellular role of galectin-9 in organizing HLA-DR at the IS appears to be independent of any putative function for gal9 in antigen processing

ICAM-3 binding to LFA-1 on mature DCs induces MHC-II clustering at the IS in a Rac1-mediated process, which in turn regulates actin cytoskeleton reorganization (de la Fuente et al., 2005). Interestingly, we have previously shown that gal9 modulates Rac1 activity to control cortical membrane structure and maintain plasma membrane integrity (Querol Cano et al., 2019). It is thus plausible to hypothesize that gal9 may influence MHC-II organization at the IS by connecting MHC-II to ICAM-3 and Rac1 signaling pathways. On the other hand, gal9 may regulate lipid raft dynamics and MHC-II trafficking by interacting with Flotillin-1 (FLOT1) and Flotillin-2 (FLOT2), key components of cholesterol-dependent membrane domains. FLOT1 and FLOT2 contribute to the sorting, trafficking, and organization of proteins critical for antigen presentation and as integral parts of lipid rafts, they serve as platforms where MHC-II complexes and TCR are clustered, facilitating effective antigen screening and presentation at the IS (Hiltbold et al., 2003; Khandelwal and Roche, 2010; Poloso et al., 2004). Our mass-spectrometry data identified FLOT1 and FLOT2 as putative gal9 binding partners and gal9 may play a role in stabilizing or modulating these cholesterol-rich membrane domains (Tanikawa et al., 2008). Further supporting this hypothesis, our RNA-seq pathway enrichment analysis revealed cholesterol-related pathways to be altered in gal9-deficient DCs, suggesting that gal9 may influence lipid raft-mediated organization of MHC-II at the IS.

In summary, this study is the first to describe an intracellular function for gal9 in organizing MHC-II recruitment and localisation at the IS, providing new insights into how galectins regulate immune receptor positioning to enhance T cell activation. Data presented here highlights a novel role for gal9 in promoting DC function, underscoring the broad impact of galectins in regulating immune responses.

## Acknowledgements

The authors thank the Radboudumc Microscopy Imaging Technology Center for the use of their microscopy facilities and Dr. Koen van den Dries for his help in analysing live cell imaging data. We thank Pascal W. T. C. Jansen and Dr Cornelia G Spruijt for their assistance with the proteomics experiments and Dr Joost Martens for his help performing RNAseq analysis. We also thank Floris van Dalen for synthesizing the BMV109 probe. The MV151 was a kind donation from Prof. Hermen Overkleeft. This work is supported by the Dutch Cancer Society (grants 11618 and 12949) and Radboud University Medical Center. ABvS is supported by the Netherlands Organization for Scientific Research (NWO), the Institute of Chemical Immunology (project ICI 000-23), ZonMW (project 09120012010023), and the European Research Council, Proof-of-Concept Grant (project 101112687).

## Data availability

All data are available upon request. Mass spectrometry dataset is deposited in the public proteomics identifications database PRIDE (dataset identifier PXD039781). All sequencing data have been submitted to GEO under accession number GSE282411.

---

**Supplementary Figure 1.**
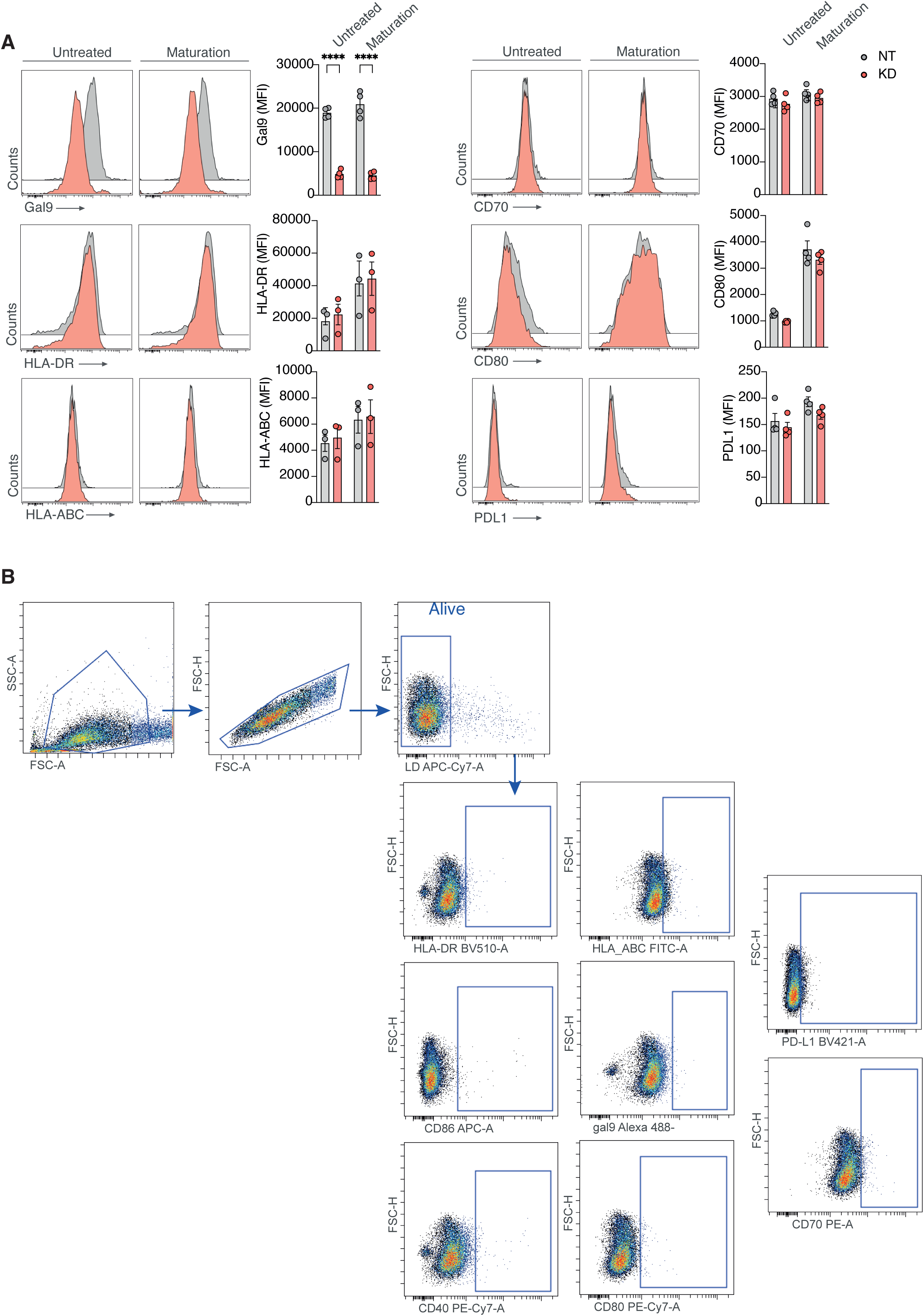
Human DCs depleted for gal9 are not impaired in their maturation. **(A)** WT (grey) or KD gal9 (red) DCs were matured with IL-1β, IL-6, PGE2 and TNFα during 18-24h and membrane expression of HLA-DR, CD70, CD80, HLA-ABC, PDL1 was assessed by flow cytometry. Representative flow cytometry plots from data shown in the graphs from one donor. Graph represents the mean fluorescence intensity (MFI) ± SEM from four donors. Two-way ANOVA with Šídák’s multiple comparisons test was conducted between conditions. ****p < 0.0001. **(B)** Gating strategy of markers in (A). Dot plots shown here are from the FMO staining controls.

**Supplementary Figure 2.**
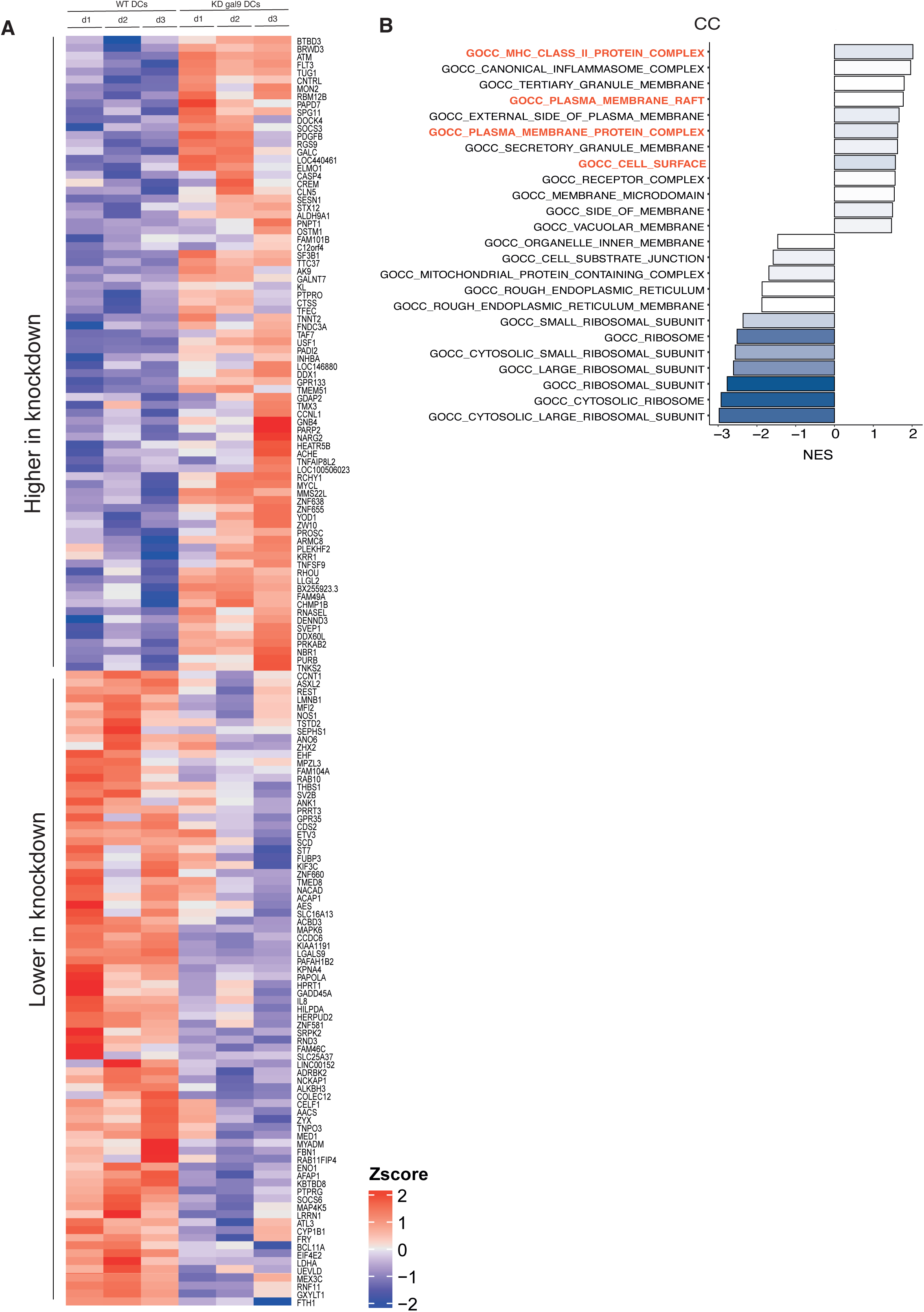
MHC-II complex assembly and plasma membrane complex formation are differentially enriched between gal9 KD and WT DCs. **(A)** Heatmap showing Z-transformed gene expression values for all genes altered in the gal9 KD cells compared to WT controls. **(B)** Results of gene set enrichment analysis using gene ontology (GO) terms, from the Cellular Compartments (CC) dataset. Significance is represented by the -log adjusted p-value. Pathways of interest are highlighted in red.

**Supplementary Figure 3.**
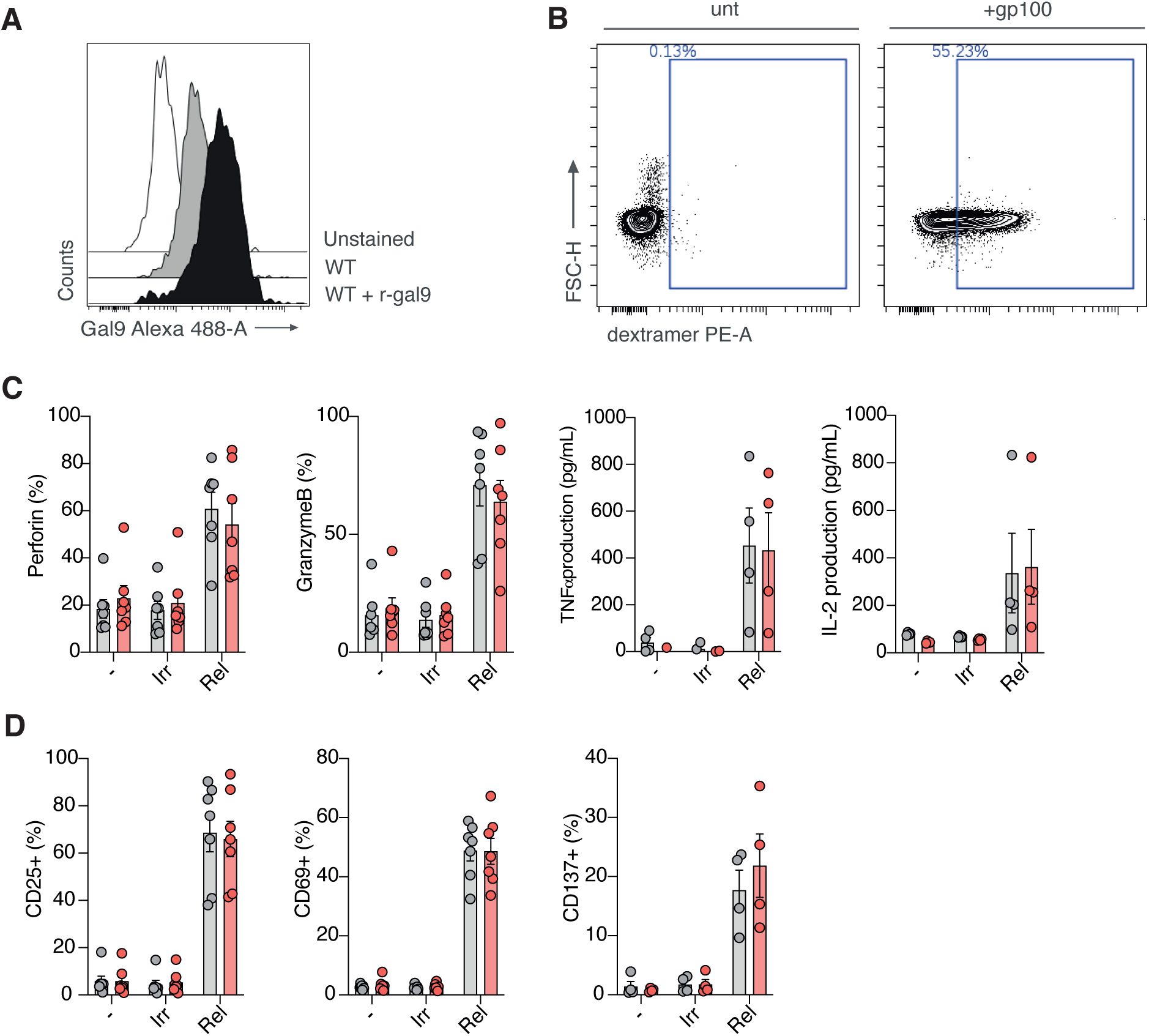
Gal9-deficient-DCs do not alter CD8^+^ T cell activation and cytotoxic capacity. **(A)** Flow cytometry histograms depicting gal9 mean fluorescence intensity in unstained (white), WT (gray) and recombinant gal9 treated WT (black) DCs. **(B)** Dot plots depicting the gp100 dextramer staining after 5h for human primary CD8^+^ T cells that were untransfected or transfected with a gp100-specific TCR encoding mRNA. **(C)** Quantification of 4-7 independent donors of different CD8^+^ T cell effector readouts for the gp100-dependent assay (perforin, granzyme B, TNFα and IL-2 production, from left to right). **(D)** Quantification of 4-7 independent donors of different activation surface markers for CD8^+^ T cells for the gp100 assay (CD25, CD69 and CD137 from left to right). Gray: T cells cultured with WT DCs. Red: T cells cultured with KD gal9 DCs. Data represent mean correlation coefficient ± SEM of three independent donors. Two-way ANOVA with Šídák’s multiple comparisons test or paired t-test was conducted between conditions.

**Supplementary Figure 4.**
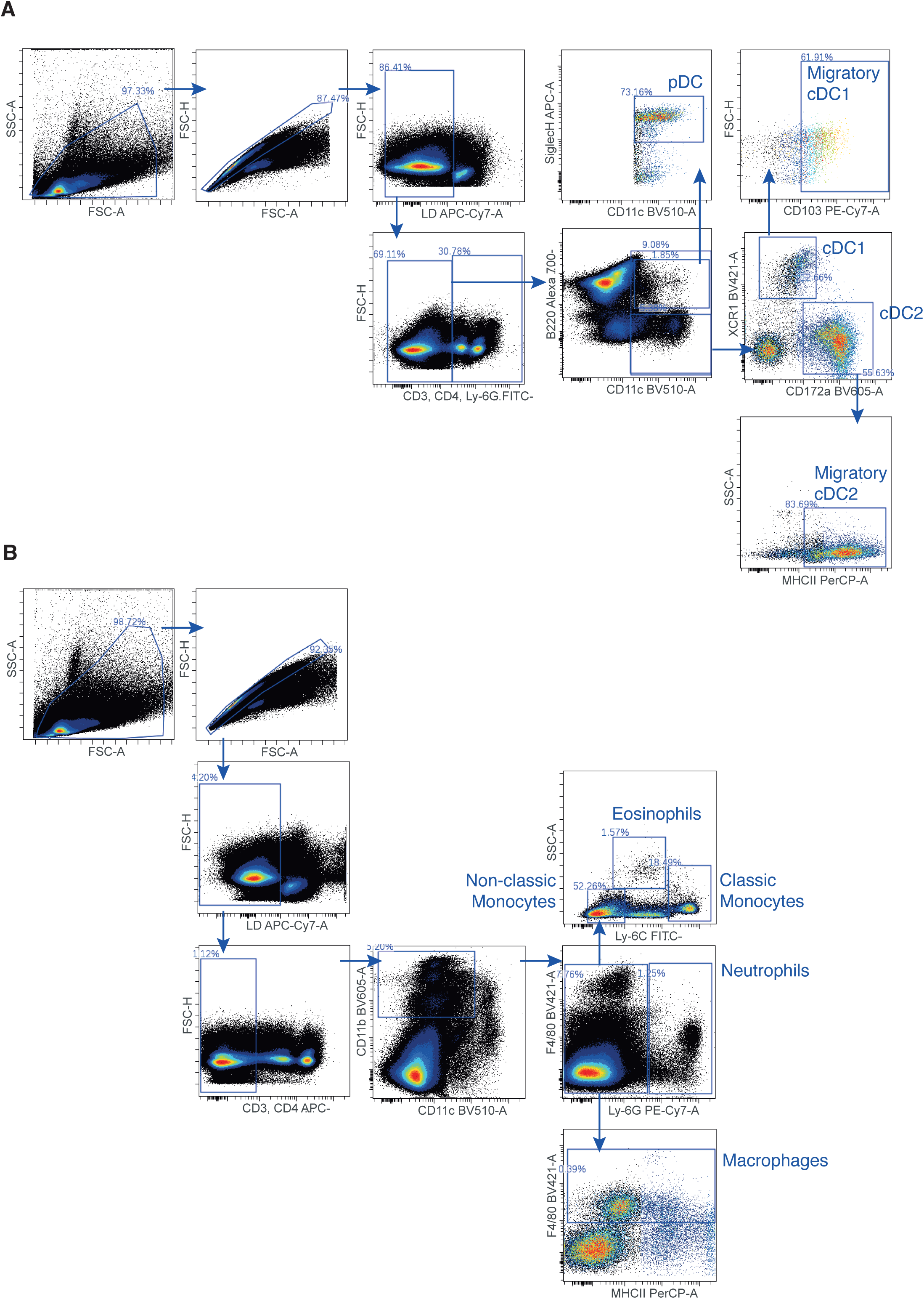
Flow cytometry gating strategy for DC subsets and myeloid populations in naïve-treated animals.

**Supplementary Figure 5.**
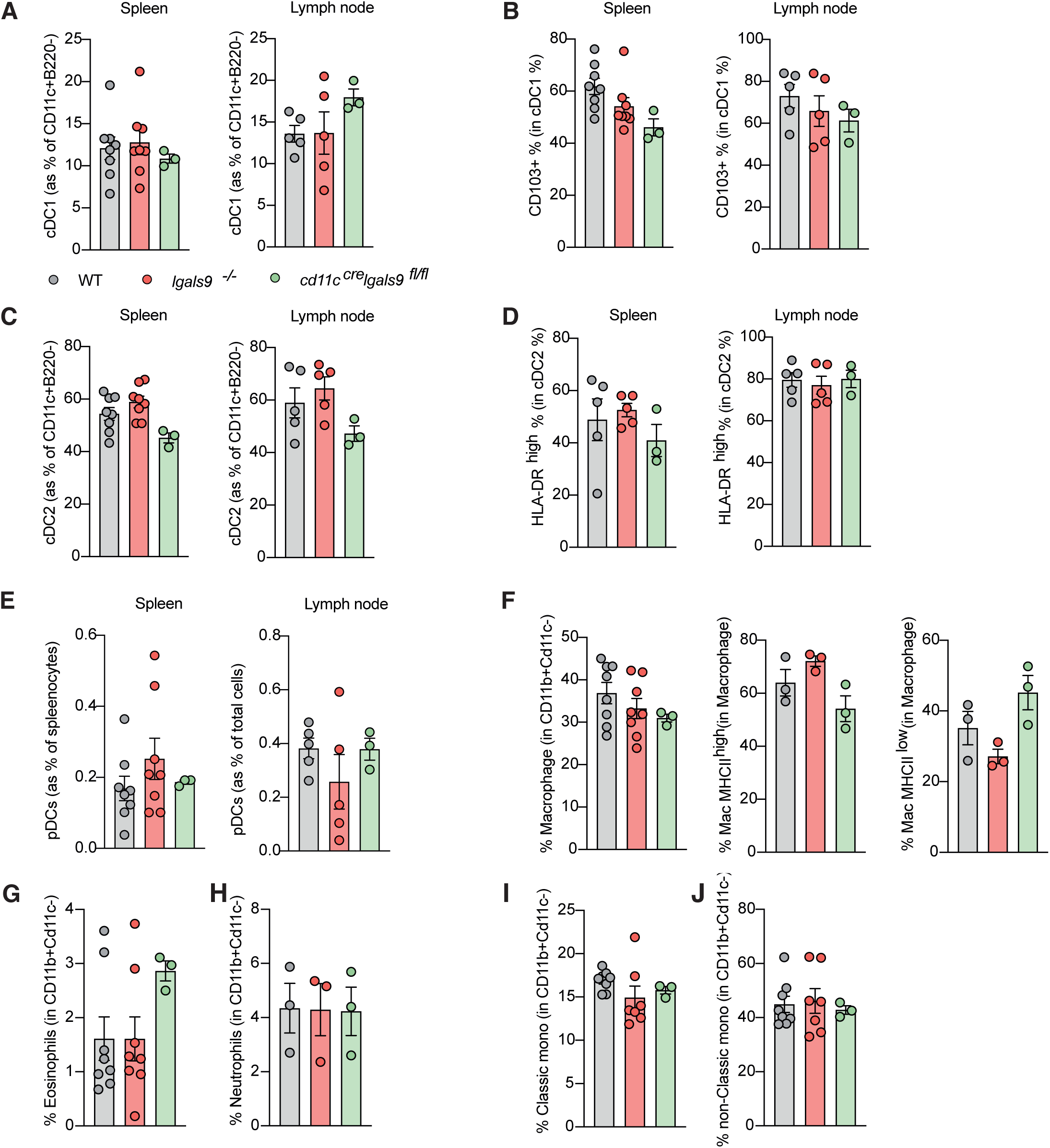
Dendritic cell development is not affected by galectin-9 / galectin-9 does not influence myeloid and DC development. **(A - J)** Percentage (%) of DC subsets and myeloid cell types in the spleen and lymph nodes of three mouse models: WT (gray), total gal9 KO (red), and conditional gal9 KO in DCs (green). Data represent mean correlation coefficient ± SEM of 3-8 independent animals. One-way ANOVA with Dunnett’s multiple comparisons test was conducted to compare conditions.

**Supplementary Figure 6.**
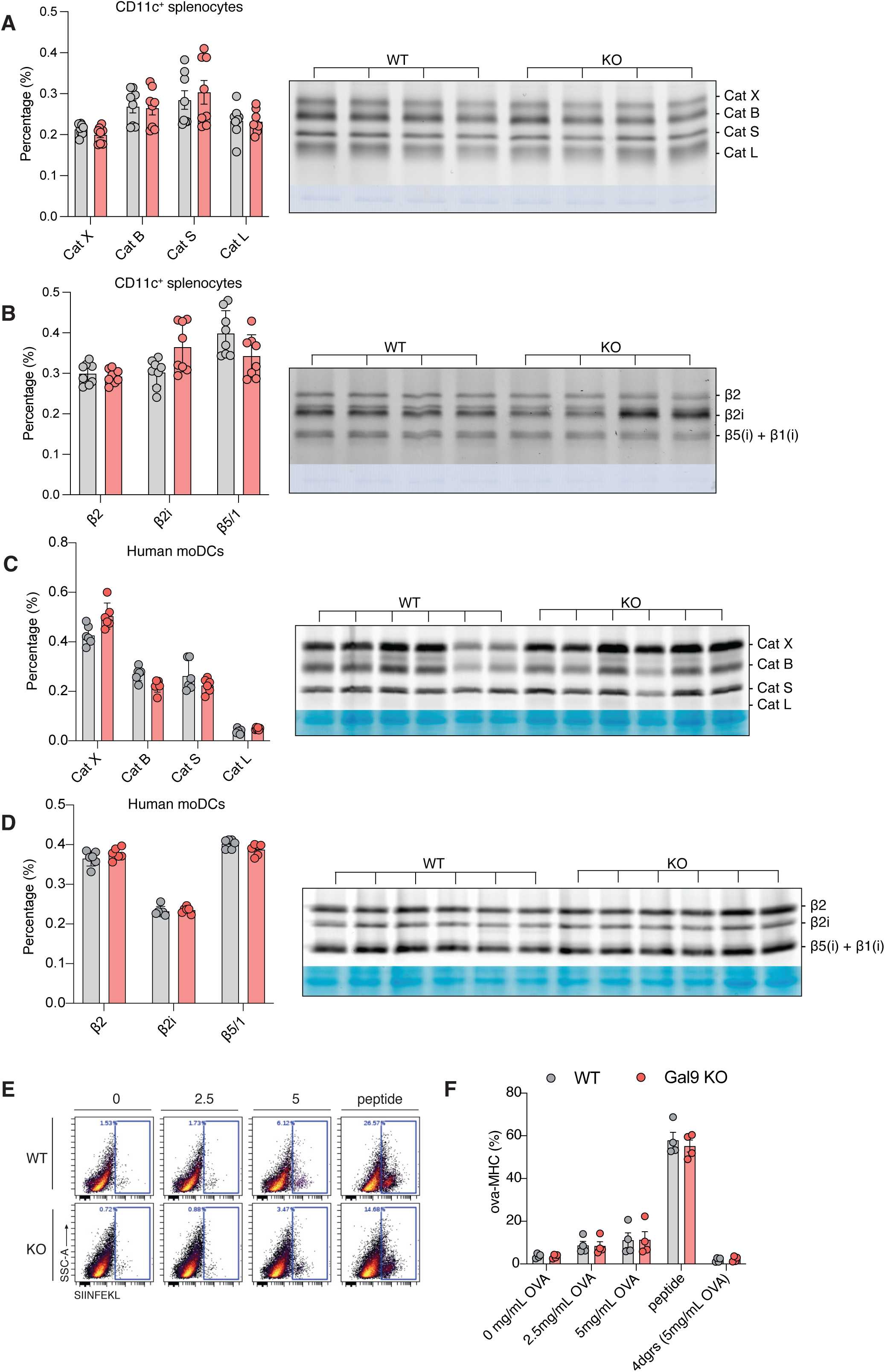
OVA-presentation, cathepsin and proteasome profiling of human and murine DCs. **(A)** Cathepsin profiling of WT and gal9 KO CD11c^+^ splenocytes. Representative SDS-PAGE is shown with cathepsins assigned. Data is shown as mean ± SD for four independent mice (technical duplicates). **(B)** Proteasome profiling of WT and gal9 KO CD11c^+^ splenocytes. Representative SDS-PAGE is shown with proteasome subunits assigned. Data is shown as mean ± SD for four independent mice (technical duplicates). **(C)** Cathepsin profiling of WT or KD gal9 monocyte-derived dendritic cells (moDCs). SDS-PAGE is shown with cathepsins assigned. Data is shown as mean ± SD for three independent donors (technical duplicates). **(D)** Proteasome profiling of WT or KD gal9 moDCs. SDS-PAGE is shown with proteasome subunits assigned. Data is shown as mean ± SD for three independent donors (technical duplicates). **(E)** Dot plot showing fluorescence intensity of the SIINFEKL peptide-bound to MHC-I detected in WT and gal9 KO murine DCs after culturing them with 0, 2.5 or 5 mg/mL OVA (or peptide as positive control) at 37° C for 18 h hours. **(F)** Quantification of data shown in (E) from four independent mice. WT (grey) or KD/KO gal9 (red) DCs. Data shown as mean ± SEM. Two-way ANOVA with Šídák’s multiple comparisons test was conducted between conditions.

**Supplementary Figure 7.**
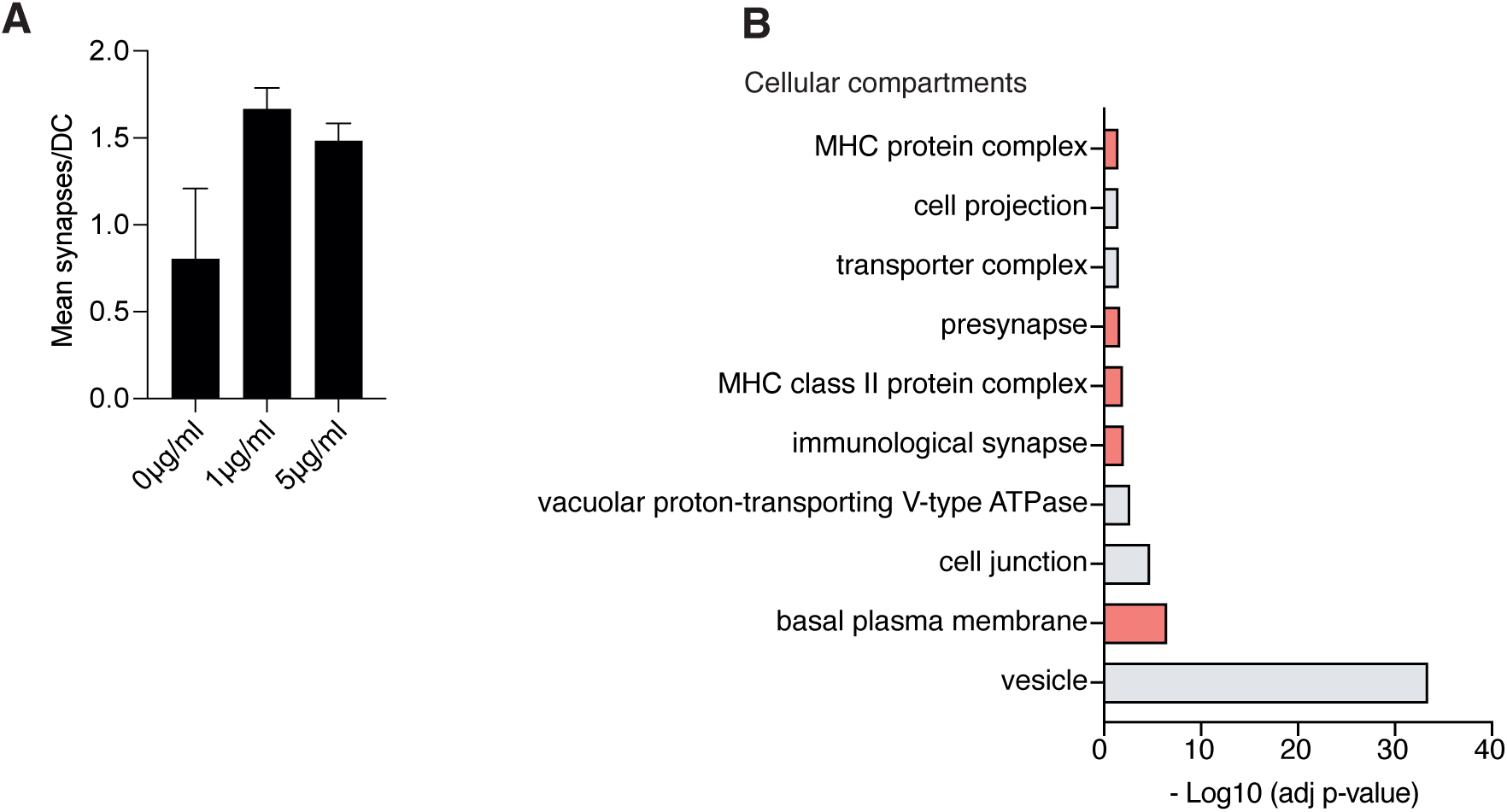
Gal9 co-immunoprecipitated proteins associate with MHC-II complex assembly and immune synapse pathways. **(A)** WT DCs were pre-incubated with 0, 1, or 5 μg/mL of superantigen SEB before being co-cultured with allogeneic T cells for 2 h. Data show number of stable DC-T cell contacts formed after this time. **(B)** Results of gene set enrichment analysis using gene ontology (GO) terms, from the Cellular Components (CC) dataset. Pathways of interest are highlighted in red. Statistical significance is represented by the adjusted p-value (-log10) (x axis).

**Supplementary movie 1.** Representative time-lapse of WT DCs co-cultured with T cells. DCs are shown in magenta and T cells in green. Scale bar = 10 µm.

**Supplementary movie 2.** Representative time-lapse of KD gal9 DCs co-cultured with T cells. DCs are shown in magenta and T cells in green. Scale bar = 10 µm.

